# Socioeconomics drive population change in the world’s largest carnivores

**DOI:** 10.1101/2021.11.30.470341

**Authors:** Thomas F. Johnson, Nick J. B. Isaac, Agustin Paviolo, Manuela González-Suárez

## Abstract

Land-use^1,2^ and climate change^3–5^ have been linked to wildlife population declines, but the role of socioeconomic factors in driving declines, and promoting population recoveries, remains relatively unexplored despite its likely importance. Here, we evaluate a comprehensive array of potential drivers of population changes observed in some of the world’s most charismatic species – large mammalian carnivores. Our results reveal a strong role of human socioeconomic development, which we find has a greater impact on population change than habitat loss and climate change. Increases in socioeconomic development are linked to sharp population declines but, importantly, once development is high, carnivore populations have the potential to recover. These links between human development and wildlife population health highlight the challenges ahead to achieve the different UN Sustainable development goals.

## Main Text

Humans are transforming the planet, driving species to extinction and altering ecosystems^6^ - pushing biodiversity closer to its planetary limit^7^. However, whilst extinction rates suggest biodiversity is extremely threatened, there is a lack of consensus across the biodiversity change literature, with recent work showing that many populations are not declining ^8–10^. Instead, environmental change causes variations in wildlife population trends across ecological communities, with some populations declining whilst others prosper ^11^; reshaping community structure and possibly altering ecosystem functions ^12,13^. One of the core challenges in understanding biodiversity change is moving beyond documenting species declines, towards models that can predict trends under environmental change to facilitate the planning and effective implementation of conservation management measures. This is not an easy task as observed population trends are not always representative of actual trends ^14,15^, trends can also be difficult to derive ^16^, and the factors that influence population trends are numerous and hard to measure ^17^. This, in-part, explains the low predictive accuracy of macro-scale biodiversity change models ^3,4^.

Previous work has focussed on understanding how land-use change ^1,2^, forest-loss ^10^ and climate change ^3–5^ influences biodiversity trends, either separately or in combination. However, focussing primarily on environmental change ignores the role of socioeconomic factors in mitigating or magnifying impacts. For example, populations are more likely to increase in areas with strong governance ^18^, and population declines can be exacerbated by armed conflicts and species life-history traits ^19^. To effectively detect the signal of environmental change impacts, it is important to consider the multidimensional context and diversity of potential factors that influence population dynamics. Here, we take this comprehensive approach to explore how land-use change, climate change, and governance impact population trends of large terrestrial carnivores globally. We also explore how species life-history traits can mitigate or magnify these trends, specifically focussing on species from the families Canidae, Felidae, Hyaenidae and Ursidae of the order Carnivora, which include the largest terrestrial carnivores on the planet.

Large carnivores (such as lions, tigers, and wolves) are an important focal group to study as they are culturally important ^20^, essential for regulating ecosystem function ^21^, and can act as indicator species of the overall status of biodiversity within an area ^22^. Large carnivores are also generally well-studied taxa with abundant trend datasets available from the primary literature across a wide spectrum of environmental change and governance scenarios. The morphology, ecology and behaviour of these taxa is also generally well described ^23^, allowing us to evaluate important influences without being impacted by poor inference from missing data ^24^. Finally, despite being well-studied, the population status of large carnivores is unclear, with reports of devastating declines ^21^ contrasted with remarkable recoveries ^25^. Studying these differences in responses can provide insight beyond these taxa, revealing strategies and scenarios that could help bend the biodiversity curve ^26^.

### Factors influencing population change

To determine how land-use change, climate change, and governance influence population trends in large carnivores, we compiled two types of trend data: 1) quantitative estimates of change which we converted into annual rates of change (%) in abundance, and 2) qualitative descriptions of change: Increase, Stable, and Decrease. By including qualitative records, we increased the sample size and importantly, the taxonomic and spatial representativeness of the trend data (Figure 1). In total, we compiled trends for 1,127 populations, sourced from 7,352 abundance estimates available in the CaPTrends ^27^ and Living Planet databases ^28^. Rates of change were available for 50 of 87 species in our focal group, with locations representing 75 countries located in all continents with native *Carnivora* species, and variable time periods between 1970 – 2015.

**Figure 1.**
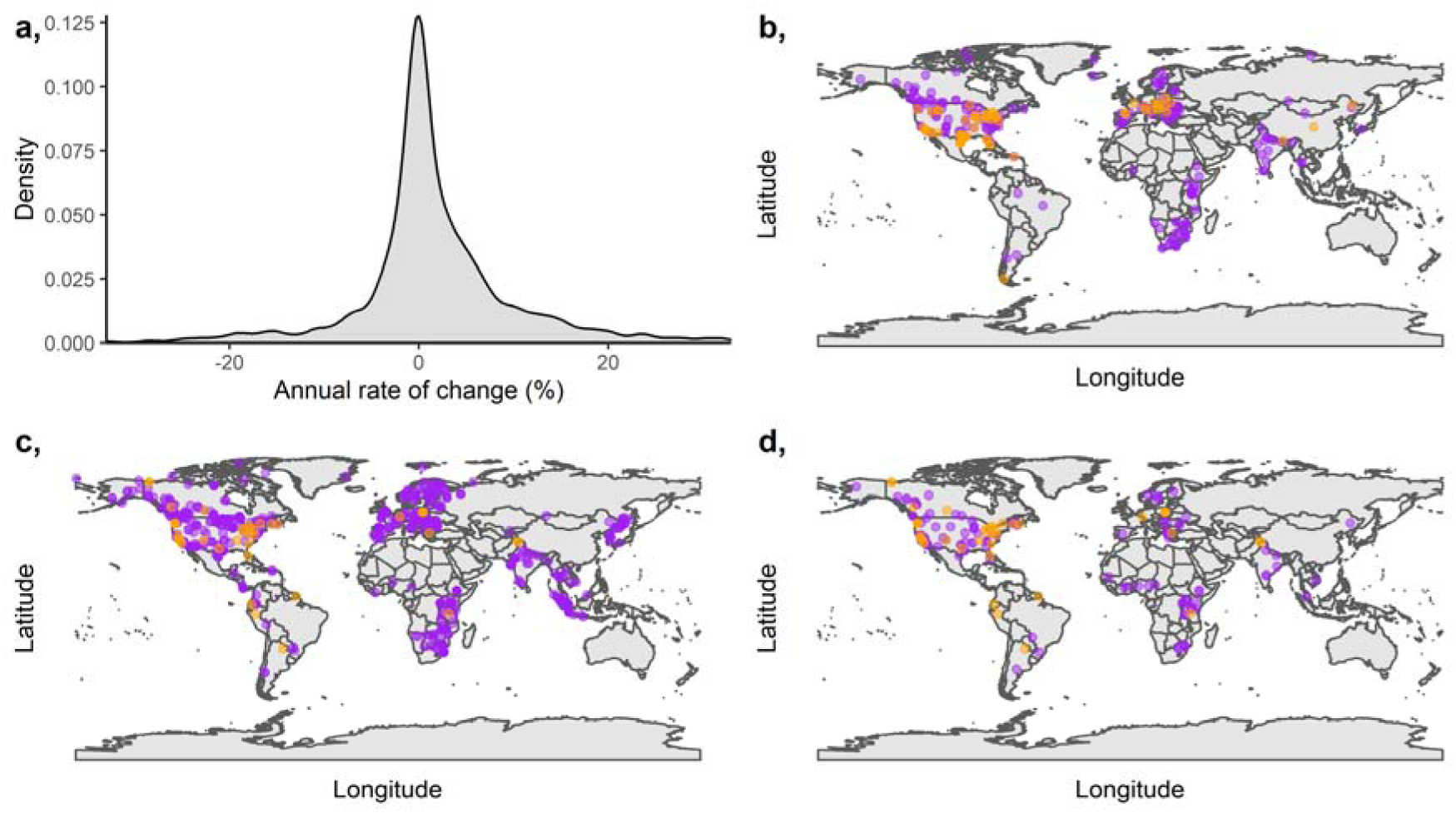
Distribution of trend data. a) Distribution of annual rates of change from quantitative trend values. b-d) Quantitative (purple) and qualitative (orange) records distributed through space, split into three groups: Increasing (b), Stable (c) and Decreasing (d). For quantitative trends, ‘Increase’ describes records with an annual rate of change exceeding 5%, ‘Stable’ range from -5% to 5%, and ‘Decrease’ trends are below 5%. Maps are plotted on a WGS84 projection.

For each trend, we estimated several metrics describing land-use, climate, and governance features, each of which we hypothesized could influence population trends (Figure 2). Using the trend data (response) and associated covariates, we developed a hierarchical Bayesian linear model (see Materials and methods). To allow the inclusion of both types of trend data within one model, which reduced taxonomic and spatial bias, we included a novel censored response-term to treat the qualitative descriptions (type 2) as partially known quantitative trends. Models included sixteen covariates, as well as six interactions specified because we hypothesized the impacts of environmental change could depend on species traits ^29^, the quality of national-level governance ^18^, and the interactions between different types of environmental change. For example, specialist species are likely to experience greater declines under combined land-use and climate change ^30^.

**Figure 2.**
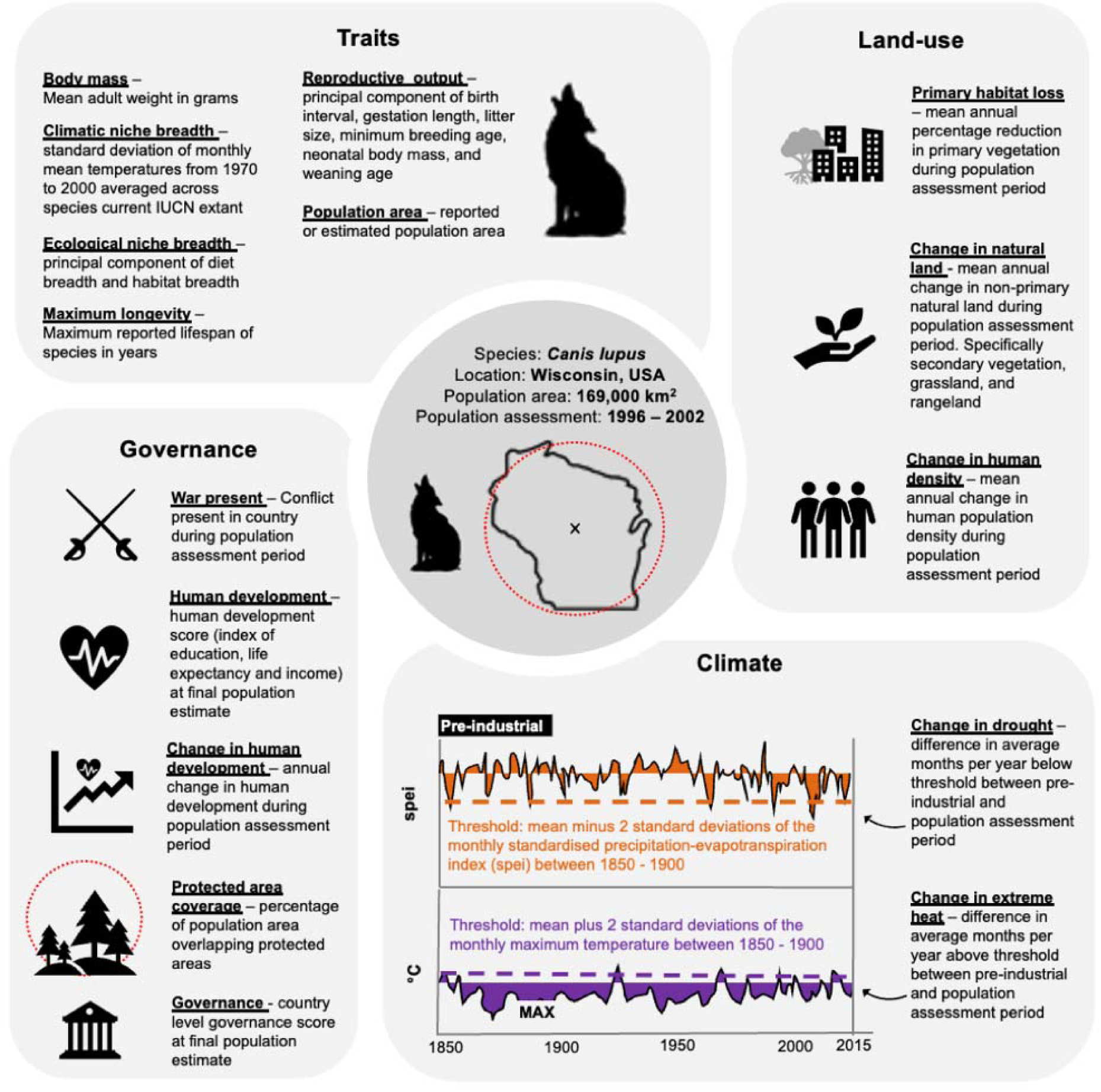
Sixteen covariates with a proposed effect on carnivore population trends. Covariates are highlighted in bold and underlined. Covariates fall in four groups: Traits, Land-use, Climate, and Governance Text alongside covariates briefly explains how the variable was derived, whilst full explanation and justifications for inclusion are available in Supplementary methods: Covariates.

#### Land-use

As predicted and shown in other taxa ^1^, we found that primary habitat loss is associated with declines in carnivores. However, we expected carnivore declines to be more extreme when habitat was associated with an increase in human density, relatively mitigated when replaced by semi-natural land, and highly dependent on the species ecological niche breadth (its degree of specialization), but we found no evidence supporting these interactions (Figure 3). Our findings suggest that all populations decline in the immediate aftermath of primary habitat loss, regardless of the species ecological specialization and what replaces the primary habitat.

**Figure 3.**
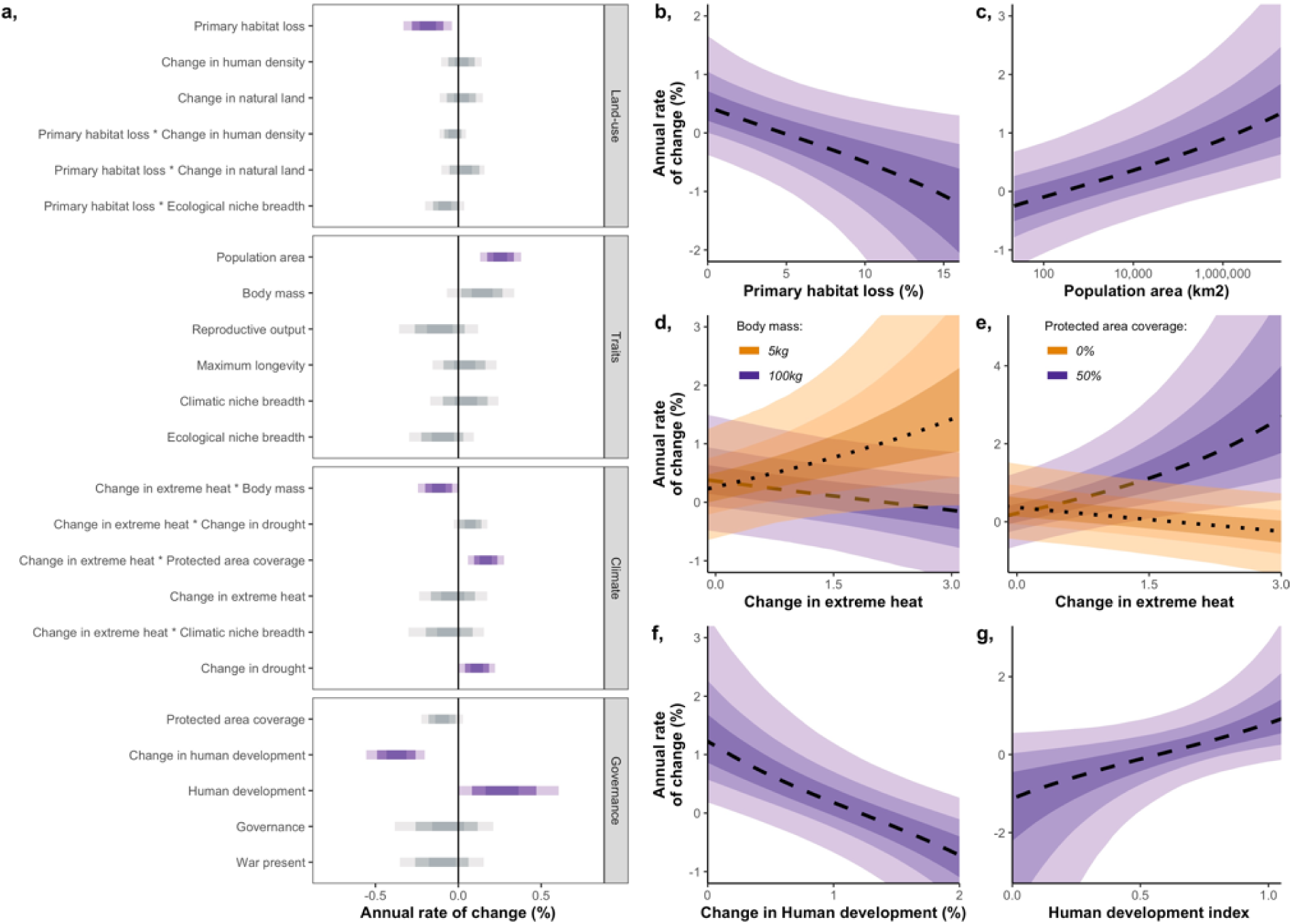
Population trend model coefficients and marginal effects. a) Annual rate of change coefficients from fixed effect parameters in a hierarchical Bayesian linear model, with 50%, 80%, and 95% credible intervals. Coefficients with an effect at the 90% credible interval are coloured in purple. b-g) Marginal effects for a selection of important covariates against median predicted annual rate of change: mean annual primary habitat loss (b); area of population buffer zone on the log_10_ scale (c); mean number of months per year where the average temperature of the population monitoring period exceeded the mean plus 2 standard deviations of the average temperature of the baseline pre-industrial period (1850 – 1900), interacting with body mass (d) and protected area coverage (e); annual change in human development over the population monitoring period (f); and human development at the final year of population monitoring (g). All covariates were back scaled from any transformations. Error ribbons represent the 50%, 80%, and 95% credible intervals.

#### Climate change

Unlike land-use, climate change effects were complex (Figure 3). First, an increasing frequency of extreme heat events had no consistent effect on carnivore trends, instead, its effect varied across species of different sizes, and populations located in areas with varying protected area coverage. In particular, extreme heat was associated with population declines in large species but with increases in smaller carnivores. Prior work has suggested large-bodied birds are susceptible to rising temperatures from climate change ^31^, and if this result holds true for other vertebrates, it may further threaten large flagship megafauna (e.g. polar bear, tiger and lion). We also found that extreme heat was associated with declines in populations outside of protected areas but rapid population growth inside, suggesting protected areas mitigate climatic extremes. As protected areas are amongst the least impacted fragments of land on the planet, they may naturally offer features that buffer extreme temperatures e.g. micro-climates from canopy cover ^32,33^. Increased population densities could also reflect increased use and movement towards these protected areas from less suitable habitat in the short term. The expansion of protected areas could be an effective approach to support future carnivore populations in the face of climate change, and likely could benefit other taxa including birds ^34^. However, protected areas cannot promote continuous population growth in the long term, as their area and resources are finite. In fact, our results show that protected areas (in isolation) have no effect on population trends. Our results reveal another interesting effect of climate change: more drought events were associated with population increases. This may seem counter-intuitive, but in the short-term it is likely that drought would lead to an unusual concentration of vulnerable prey around the remaining water sources, making their encounter and capture, both predictable and easy for predators. The increase in food availability for carnivores could explain this population growth.

However, such benefits of drought would likely be short-term, as predator-prey dynamics would eventually cause carnivore population declines. Our analyses reveal interesting and complex relationships between climate variables, species traits, and habitat protection that should be further investigated in other taxa and monitored over time to detect and respond to changes.

#### Governance

Carnivore populations were more likely to increase in areas with high human development scores (Figure 3). Human characteristics associated with development (i.e. improved quality of human life) appear to be more important for large carnivores than higher-level governance characteristics (i.e. rule of law and legislation). These findings differ from previous work showing bird declines are greatest in areas with weak governance ^18^ and attributed the recoveries of Europe’s carnivores to the strong governance of the European Union ^25^. Wildlife can receive regulatory protection through governance, but perhaps protection may also be achieved by improving human quality of life, and in turn, tolerance for wildlife. For example, if a carnivore kills the livestock of an owner living in extreme poverty (low human development), there is greater potential for human-wildlife conflict regardless of the species regulatory protection.

Whilst carnivore population increases were most likely in areas with high human development, there was a cost to reaching this development level, as we found rapid human development growth was associated with carnivore population declines (Figure 3). This relationship does not appear to be driven by underlying factors stimulating human development growth (e.g. detrimental land-use changes, or natural resource extraction for economic growth), as these factors were directly modelled and had a less clear statistical association despite being measured at more relevant spatial-scales. Instead, we hypothesize two potential mechanisms to explain this result. First, areas with slow human development growth (global north) generally correspond to countries where carnivore populations were threatened or extinct before systematic carnivore data collection began, e.g. the grey wolf was extirpated from the UK more than 500 years ago. As such, these countries likely have a lower baseline carnivore population and greater capacity for recovery, or have already lost the carnivore species that are most susceptible to human impacts. However, a lower baseline cannot explain the association between fast human development growth and carnivore population declines, for this association, we hypothesise that human development change provides a more holistic snapshot of environmental and societal transformation, including underlying factors as well as changes in culture and relationships with wildlife. For example, development in Kenya has seen increases in urbanisation and infrastructure as well as changes in people’s relationships with nature ^35^.

#### Relative contributions

To further assess the contribution of important covariates, we developed three counterfactual scenarios which describe how the absence of specific features (or threats) would alter the 1,127 observed trends. Scenarios included: 1) No loss in primary habitat – primary habitat loss set to zero; 2) No climate change – both change in frequency of extreme heat and drought set to zero; and 3) No growth in human development – change in human development set to zero. We subtracted these counterfactual predictions from the observed trends to define ‘Difference in annual rate of change (%)’, whereby a positive value indicates carnivore populations would be in better shape (fewer declines) had there been no changes in habitat, climate and human development, whilst a negative value indicates observed changes benefitted carnivore populations (more declines under no change).

The counterfactual scenarios reveal the great importance of human development as its contribution to population changes dwarfed those of land use and climate change (Figure 4). No consistent increases or decreases in population trends resulted from assuming observed habitat loss and climate change had not occurred (although African carnivore populations would have overall benefitted from preservation of habitat). However, human development changes had clear and consistent impacts across carnivore populations in all continents, with nearly all counterfactual trends being higher.

**Figure 4.**
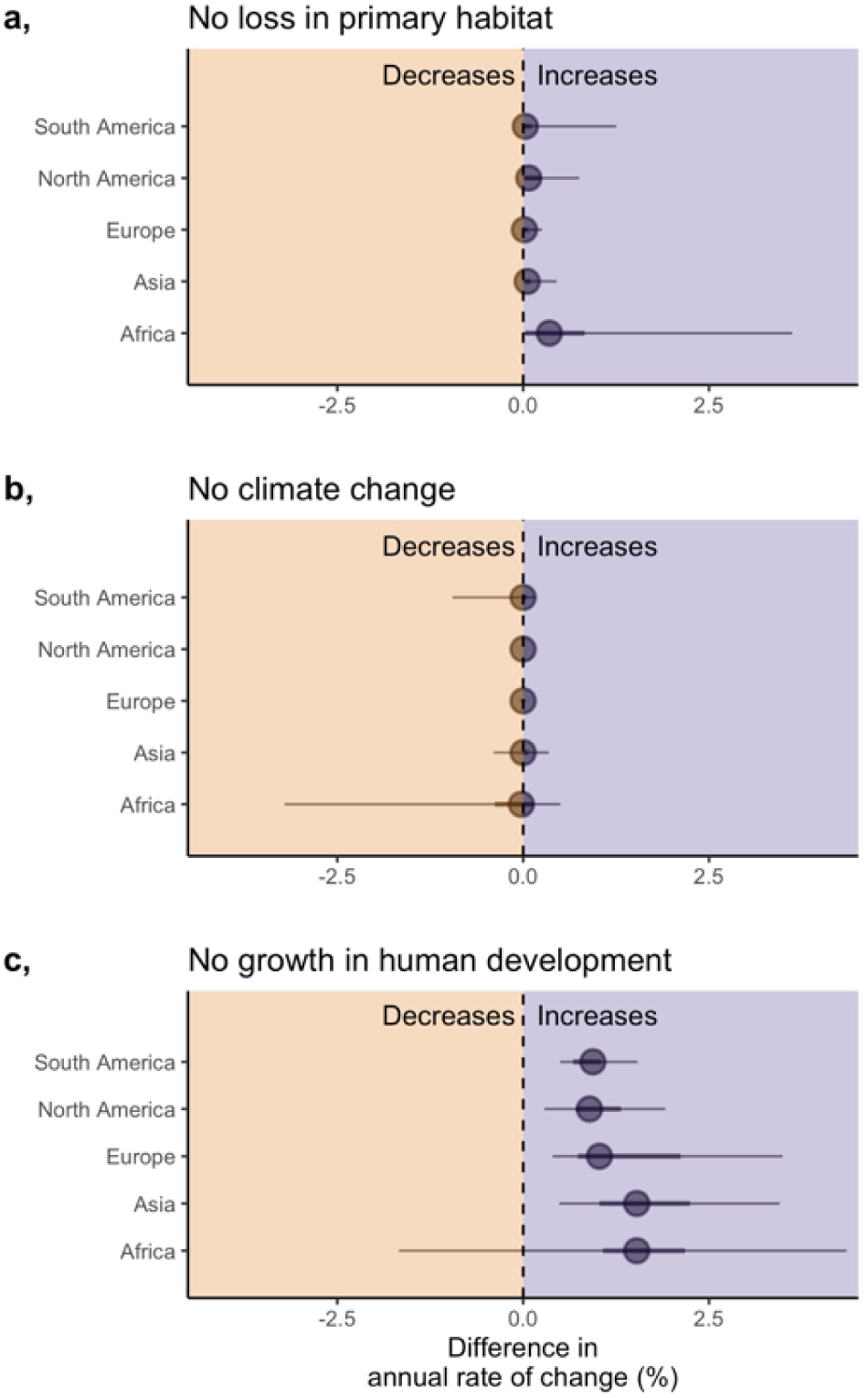
Counterfactual scenarios describing the difference in the annual rate of change (%) across the 1,127 studied populations had there been no primary land loss (a), climate change (b), or growth in human development (c). Points falling on the right of the dotted line indicate that the population would be predicted to increase had observed land loss, climate change, and human development growth not occurred, and populations on the left of the line would be predicted to decrease. Points represent the median difference in the annual rate of change (predicted trend using counterfactual data minus the predicted trend using the observed data), with 50% and 95% quantiles.

While we cannot establish the direct mechanisms by which rapid human development can lead to population declines, our results identify a potential trade-off between healthy carnivore populations and human development that could hinder the achievement of the UN sustainable development goals (SDG) in developing countries. For example, improvements in health, education and income (SDGs 1-5) could negatively impact large carnivores (and biodiversity as a whole), hindering progress towards SDG 15. Work is needed to establish and mitigate the mechanisms by which human development can induce carnivore declines to resolve conflicting SDGs, whilst continuing to promote human development growth.

### Conclusions

Developing macro-scale models of population change is challenging as response data are biased ^14^ and hard to summarise ^16^, and response-covariate relationships are likely complex and numerous ^36^. Within our workflow, we attempted to address these challenges, and overall, this allowed us to achieve a moderate model fit (conditional R^2^ ∼ 0.4). We minimised biases in the trend data by integrating qualitative trends with quantitative estimates, which allowed us to increase the taxonomic and spatial scale of the work. While we could not avoid some biases, we found inference was similar across different fragments of the data and model structures (Supplementary results: Sensitivity analysis). We also attempted to capture complexity by covering a more comprehensive array of covariates than previous analyses, but we still lack data on likely important aspects that are cryptic and difficult to measure (e.g. poaching, persecution, culling, and the conservation benefits of being flagship species). There are temporal lags between disturbance-events and observable changes in the population ^10^ and we tested several to incorporate the lag that maximised model fit. However, it is possible that responses to different types of disturbance (e.g. habitat loss and climate change) have different lags, although this has not be quantified. Long lags (the maximum lag we explored was 10-years) may also occur and be associated with slow recoveries, but an absence of longer temporal extents in the response and covariate data largely prohibits this analysis at global scales (long temporal extent data is less available outside of the global north).

Our study offers a comprehensive analysis of global population trends for large mammalian carnivores. We show that abundance trends have been influenced by stressors like land-use and climate change with some interactions and effects magnified or mitigated by species traits and the protected status of the land. For climate change, we expect many of the threats are yet to be fully realised and could become more sinister in the future ^5^. These effects may not be very surprising but the particularly strong importance of human population characteristics (i.e. human development) was unexpected. Socioeconomic factors seem to be the theatre where effects of climate and land-use change are played out. We must consider the wider socio-economic scope when evaluating biodiversity changes. Focussing solely on stressors like land-use and climate change may be effective at identifying causes of declines, but provides few opportunities to identify the features that support recovery, and these features may hold the key to bending the biodiversity curve. In large carnivores, recoveries are already underway in parts of the world ^25^, but challenges remain. Capturing the multitude of complex features influencing biodiversity change will be essential to understanding and addressing these challenges.

## Acknowledgments

Thanks to Stephen Freeman for sharing code and discussing methods, as well as Chris Clements and Tom Oliver for reviewing an early version of the work.

## Data and materials availability

Data are in the process of being published and so will be open access. A link to an early draft of the CaPTrends data can be viewed at https://github.com/GitTFJ/Carnivore-population-trends-manuscript/blob/main/DataPaperV0.5.docx. Once published, all data will be publicly available. All code are available at https://github.com/GitTFJ/Carnivore-population-trends-manuscript.

## Methods

### Population trends

We sourced population trend information for species in the families Canidae, Felidae, Hyaenidae, and Ursidae of the order Carnivora from two large trend datasets: CaPTrends ^1^ and the Living Planet Database ^2^. Combined, these datasets produce a cumulative 1,127 trends (after removing duplicates and records we unreliable or unsuitable), derived from more than 10,000 population estimates. In the Living Planet Database, and for most records in CaPTrends, trends are reported as a timeseries of abundance (or density) estimates. We modelled these timeseries with log-linear regressions, where abundance (the response) was log_e_ transformed, and year of abundance estimates was selected as the predictor. We extracted the slope coefficient which represents the annual instantaneous rate of change, sometimes called the population growth rate (*r_t_*). Alongside the abundance timeseries, CaPTrends also has three other broad datatypes, all of which we converted into an annual instantaneous rate of change (*r_t_*): 1) a mean finite rate of change; 2) estimates of percentage abundance change between two points in time; and 3) timeseries’ of population change estimates (e.g. in year 1 the population doubled and in year 2 it halved). We converted all annual instantaneous rates of change into an annual rate of change percentage to improve interpretability. These annual rates of change ranged from -75% to 68%, but the majority of values fell within -10% to 10% (Figure S1a).

Alongside the quantitative records, 138 populations in the CaPTrends dataset were only described qualitatively with categories: Increase, Stable, and Decrease. These records were more common for populations located in traditionally poorer-sampled countries (e.g. with lower human development), so whilst they are less informative (only describing the direction and not the magnitude), we deem them important to reduce known biases (Figure S1b-d). As a result, we used a combination of percentage annual rates of change (N = 989) and qualitative categories (N = 138) as our response in our model – see below.

### Covariates

For each population, we extracted sixteen covariates that fell into four categories: land-use, climate, governance, and traits. Our covariates were designed to cover a diverse array of factors that could influence population trends in large carnivores. Each covariate is described briefly in Figure 2 with full descriptions of how variables were derived in the supplementary material: Covariates.

One of the challenges in identifying how covariates – which can vary in space and time - impact population trends is matching the spatial and temporal scale of the covariate with the population i.e. how much of the population is affected by the covariate at a given point in time. To tackle the spatial element of this problem, we used data on the area of extent of each population to generate a circular distribution zone around the population’s coordinate centroid. We refer to this as the ‘population area’ hereafter. We then sampled covariate values within each population area, with more sampling points in larger areas (range: 13 – 295 sampling points, Figure S3b). For covariates which varied time, we extracted the covariates across the ‘population monitoring period’, which refers to the period (start and end year) the population was monitored for. However, as evidence suggests there is lag period between a covariates impact and any detectable changes in population abundance ^3^, we tested how 0-, 5-, and 10-year lags in covariates changed model fits and effect sizes. We implemented these lags by extending the start of the population monitoring period backwards for each given lag e.g. for a 10-year lag, a normal population monitoring period of 1990-2000, would then capture covariates between 1980-2000. Sensitivity analysis showed a 10-year lag had the greatest balance of improved model fit, with high taxonomic and spatial coverage (Figure SX and Table SX.

### Modelling

At its core, our model is a linear mixed effects model, regressing annual rates of change against a combined 23 covariates and interactions, using random intercepts to account for phylogenetic and spatial nesting. The model was written in BUGS language and implemented in JAGS 4.3.0 ^4^ via R 4.0.3 ^5^. The model structure is summarized below, with a full description in Supplementary: Modelling.

#### Response

We modelled our annual rate of change response with a normal error prior. However, to allow the two different types of population trend data (quantitative rates of change and qualitative descriptions of change) to be included in the same model, we treated the qualitative records as partially known. Specifically, we censored the qualitative records to indicate that the true value is unknown, but it occurs within a specified range, with annual rate changes ranging from -50% to 0%, -5 to 5%, and 0% to 50% within the decrease, stable and increase categories, respectively. These censoring range thresholds are similar to the range of the observed rates of change (−75% to 68%). Many of the qualitative records occur in less-well represented regions, species, and time-periods, so their inclusion addresses known data biases (Figure S1). However, these lower quality records can be more prone to error. As a result, we developed a weighting term within the model to inflate uncertainty around trends derived over a short timeframe, with few abundance observations, and less robust methods – see Supplementary: Modelling – Weighted error.

#### Covariates

Prior to modelling, we identified that some of our covariates were missing values (e.g. some species were missing Maximum longevity values), which can be problematic for inference if ignored ^6^. We used imputation approaches ^7,8^ to predict these missing values and recorded the associated imputation uncertainty alongside these predictions. Within our model, we accounted for uncertainty in the imputed estimates by treating imputed values of the covariates as distributions instead of point estimates. Specifically, for each imputed value we assigned a normal distribution centered at the mean imputed estimate and with an error varying by the imputed observation standard deviation. This approach allowed us to capture imputation uncertainty and improve inference robustness.

With a combined 23 covariates and interactive effects, we were conscious of overparameterizing the model. As a result, we split these parameters into three groups: 1) core parameters – which included main effects that were considered likely drivers of population change; 2) optional parameters – which included main effects we considered interesting but with little evidence to-date of any influence on trends; and 3) interaction parameters – which includes all interaction terms between parameters. We included our core parameters (Change in human density, Primary land loss, Population area, Body mass, Change in extreme heat, Governance, and Protected area coverage) in every model, but used Kuo and Mallick variable selection ^9^ to identify important parameters from the optional and interaction groups.

#### Random intercepts

We used a hierarchical model structure to account for phylogenetic and spatial non-independence in the data, including species as a random intercept nested with genus, and country as a random intercept nested within sub-regions, as defined by the United Nations (https://www.un.org/about-us/member-states).

#### Model running

We ran the full model through three chains, each with 120,000 iterations. The first 20,000 iterations in each chain were discarded, and we only stored every 10^th^ iteration along the chain (thinning factor of 10). We opted for a large chain and burn-in due to the model complexity, and to allow a broad selection of parameter combinations to be tested under variable selection. We assessed convergence of the full model on all parameters monitored in the sensitivity analysis, as well as the model intercept, and all 23 main and interactive effect slope coefficients. We checked the standard assumptions of a mixed effect linear model (normal residuals and heterogeneity of variance), and tested the residuals to ensure no spatial (Moran’s test) or phylogenetic (Pagel’s lambda) autocorrelation. We also conducted posterior predictive checks to ensure independently simulated values were broadly reminiscent of model predicted values.

We report the median slope coefficient and associated credible intervals for each of the main and interactive effects, and produce marginal effect plots for a selection of important parameters. These marginal effects hold all other covariates at zero (which is the equivalent of the mean, as covariates were z-transformed).

### Counterfactual scenarios

To explore how observed changes in land-use, climate and human development have influenced population trends, we developed three counterfactual scenarios, where we compared observed population change to predicted population change if land-use, climate, and human development remained static. For instance, in the climate change counterfactual scenario, we predicted each observations population trend using the available covariate data (e.g. land-use, governance and trait covariates), as well as taxa and location data (to provide sensitivity to the models varying random intercepts), but set the climate change covariate data to zero (in this case, change in extreme heat and change in drought). Using this scenario data, we predicted each trend using the global model (all covariate parameters). We then subtracted these counterfactual predictions from the observed trends to define ‘Difference in annual rate of change (%)’, whereby a positive value indicates carnivore populations would be in better shape (fewer declines) under the counterfactual scenario, and vis-versa. We summarise counterfactual scenarios by reporting the median Difference in annual rate of change and 95% quantiles across the observed 1,127 populations.

## Supplementary

### Methods

#### Population trends

We sourced population trend information for all species in the families Canidae, Felidae, Hyaenidae, and Ursidae of the order Carnivora from two large trend datasets: CaPTrends ^1^ and the Living Planet Database ^2^. CaPTrends contributed 1,220 trends, and the Living Planet Database contributed 350, combining to produce a cumulative 1,474 unique (non-duplicated) trends. In the Living Planet Database, and for most records in CaPTrends, trends are reported as a timeseries of abundance (or density) estimates. We modelled these timeseries with log-linear regressions, where abundance (the response) was log_e_ transformed, and year of abundance estimates was selected as the predictor. We extracted the slope coefficient which represents the annual instantaneous rate of change, sometimes called the population growth rate (*r_t_*). There are also other formats of quantitative trends in CaPTrends which fall into three broad datatypes, all of which we converted into an annual instantaneous rate of change (*r_t_*):

1. Finite rate of change

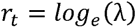 Where λ represents the mean annual finite rate of change.
2. Estimates of relative abundance change between two points in time (e.g. percentage or fold change in the past 10 years)

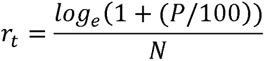 Where *P* represents the additive percentage change (e.g. a population doubling in size = 100%), and *N* is the difference in time (in years) between the two estimates of abundance. For fold changes, we first converted the fold change into an additive percentage change.
3. Timeseries of population change estimates, reported as either population lambdas or percentage changes e.g. in year 1 the population doubled (λ = 2) and in year 2 it halved (λ = 0.5). We back-converted the change estimates into abundance estimates against a constant value of 100. We then fitted log-linear regressions with abundance and year, as in the abundance timeseries.

We converted all annual instantaneous rates of change into an annual rate of change percentage to improve interpretability. These rates of change ranged from -75% to 68%, but the majority of values fell within -10% to 10% (Figure S1a).

**Figure S1.**
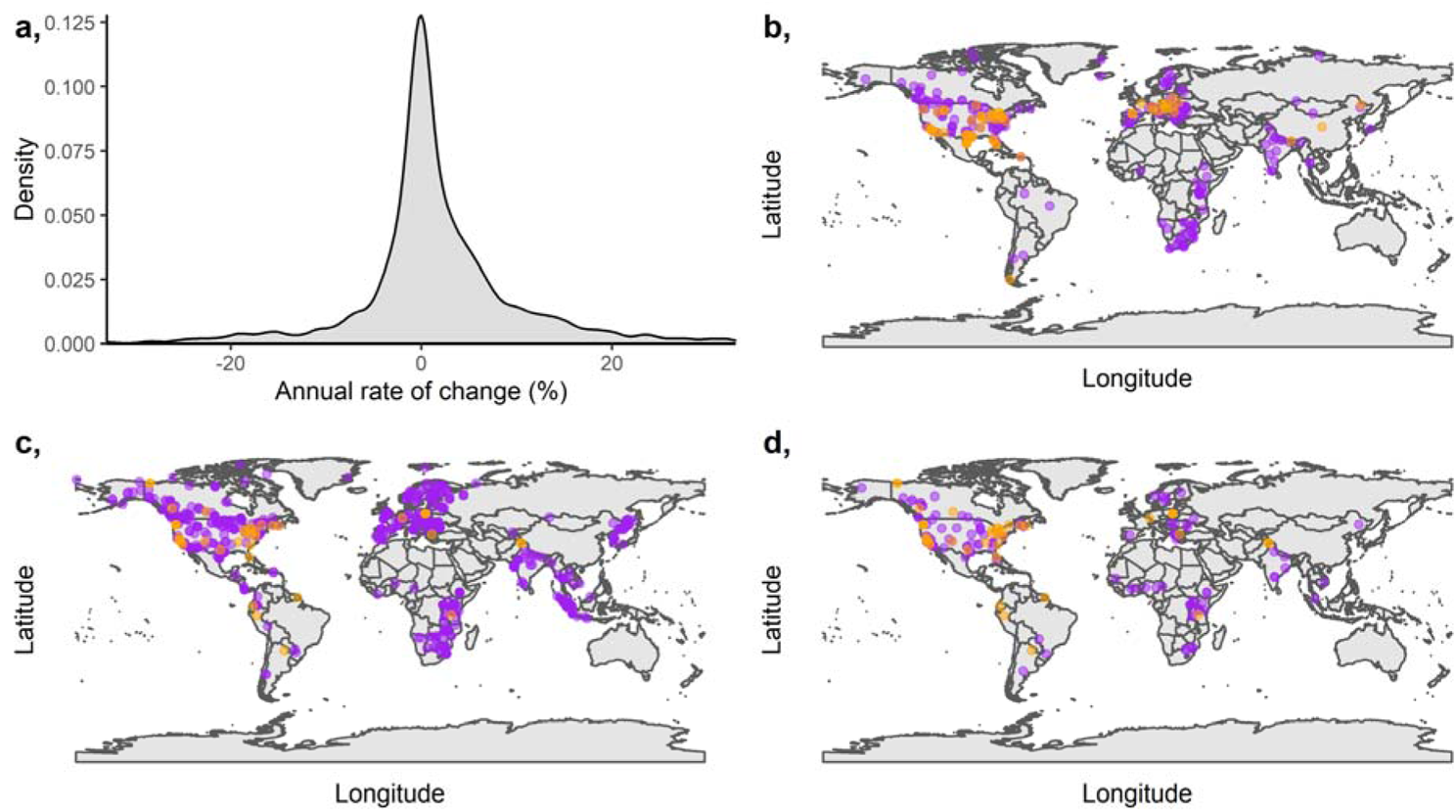
a) Distribution of quantitative records (represented as annual instantaneous rates of change); rates of change range from -75% to 68%, but axes have been trimmed to more clearly represent the bulk of the data. b-d) Spatial representation of quantitative (purple) and qualitative (orange) records, split into increasing (b), stable (c), and decreasing (d) trends. For the quantitative records, trends exceeding an annual rate of change of 5% were classed as increasing, between -5% and 5% were classed as stable, and less than 5% were classed as decreasing. The qualitative records fell naturally into the increasing, stable, and decreasing categories.

Alongside the quantitative records, 138 populations in the CaPTrends dataset were only described qualitatively with categories: Increase, Stable, and Decrease. These records were more common for populations located in traditionally poorer-sampled countries (e.g. with lower human development), so whilst they are less informative (only describing the direction and not the magnitude), we deem them important to reduce known biases (Figure S1b-d). As a result, we used a combination of annual rate of change (%) and qualitative categories as our response in our inference model – see below.

#### Covariates

Our covariates fall into four categories: land-use, climate, governance, and traits (Figure S2). One of the challenges in identifying how covariates impact population trends is matching the spatial scale of the covariate with the population i.e. how much of the population is affected by the covariate. To tackle this problem, we used data on the area of extent of each population to generate a circular distribution zone around the population’s coordinate centroid. We refer to this as the ‘population area’ hereafter. In populations without a reported extent (N = 347), we searched the locality and location description online to identify the approximate size of the population area. For example, the top result in a Google search for ‘Serengeti area’ described the location as 30,000km^2^. If we could not find an area, we assigned the population as one of the following categories: small locations (e.g. towns and counties): 1,000km^2^ [N = 123], medium locations (e.g. regions and states): 10,000km^2^ [N = 151], and large locations (e.g. countries): 100,000km^2^ [N = 73].

**Figure S2.**
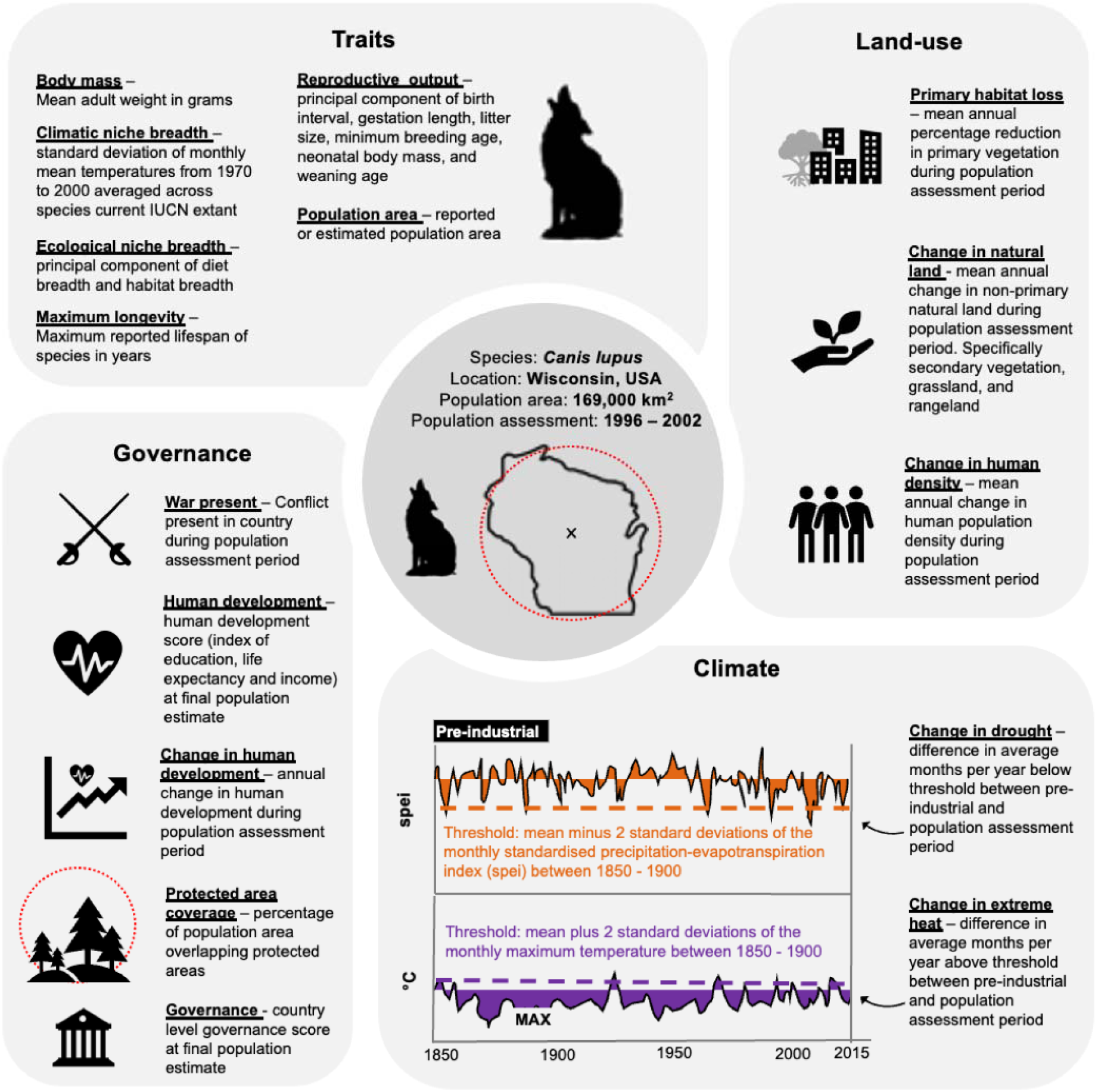
Sixteen covariates with a proposed effect on carnivore population trends highlighted in bold and underlined. Covariates fall in four groups: Traits, Land-use, Climate, and Governance. Text alongside covariates briefly explains how the variable was derived, whilst full explanation and justifications for inclusion are available in Supplementary methods: Covariates.

Given population areas regularly exceeded 10,000km^2^ (Figure S3a), it was not computationally feasible to extract covariates over the entire area; thus, we sampled from a random selection of points within each population area, sampling more frequently in larger areas (range: 13 – 295 sampling points, Figure S3b). Random sampling was only used for land-use and climate covariates, as governance covariates are measured at the national level, and all traits (except for population area itself) represent species level averages. The population areas and corresponding sampling points were defined with a Mollweide equal-area projection, but we transformed areas and points back into a WGS84 projection to match all covariate rasters (see below). In all covariates, ‘population monitoring period’ refers to the period (start and end year) the population was monitored for.

**Figure S3.**
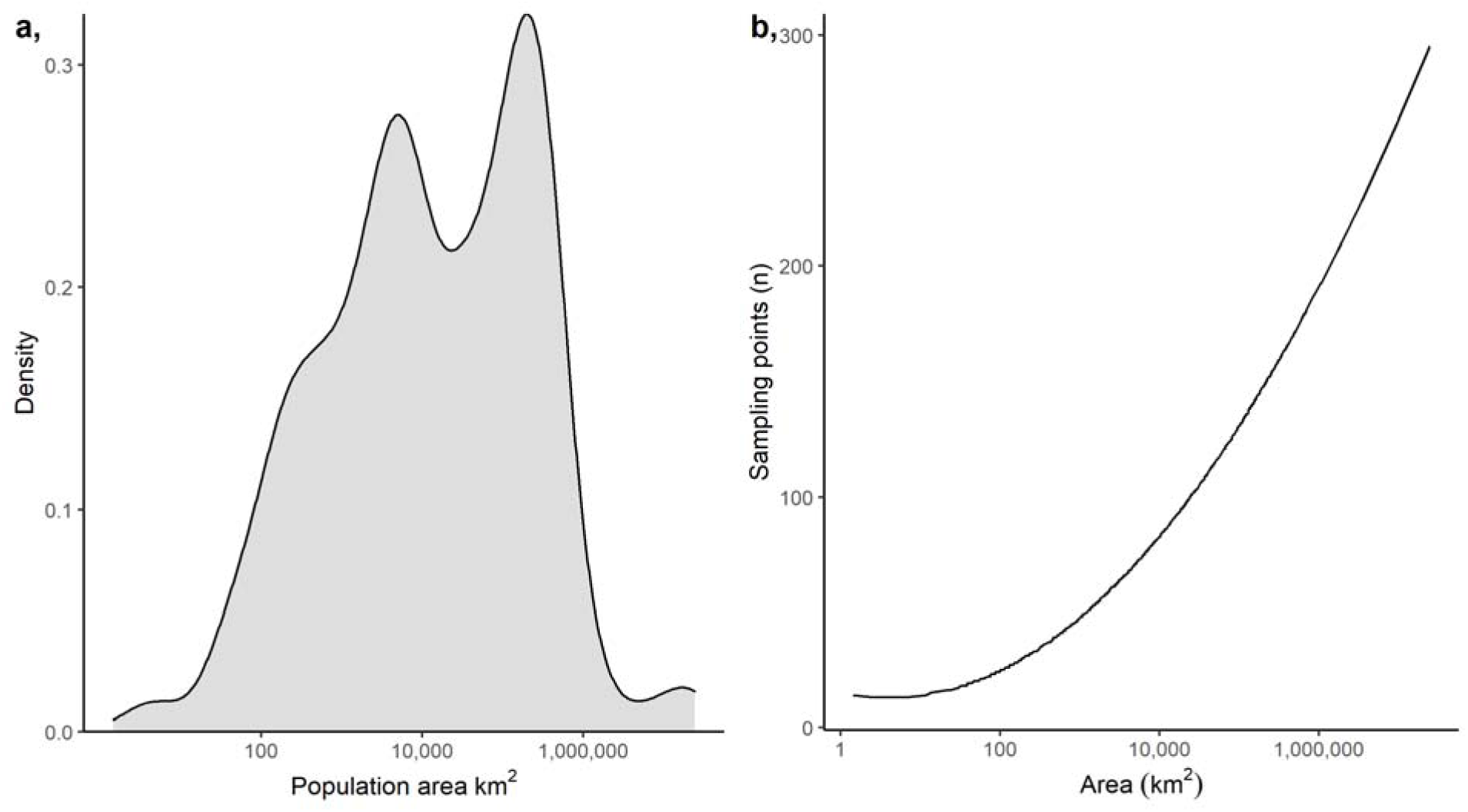
a) Distribution of population areas, the area of extent of population monitoring for 1,474 population trends extracted from CaPTrends ^1^ and the Living Planet Database ^2^. b) Frequency of covariate sampling points relative to population area size, where populations occurring over larger areas receive more covariate sampling. Both x-axes are on a log_10_ scale.

##### Land-use

We extracted three land-use covariates: Primary habitat loss, Change in natural land, and Change in human density. Primary habitat loss and Change in natural land were derived from the land-use harmonization dataset ^3^, which reports the annual proportional coverage of 11 land-use types between 1850 and 2015AD, at a 0.25° spatial resolution. To make the land-use types more biologically relevant to predators, we amalgamated a selection of the 11 types into two summary-types: primary habitat – the sum of ‘forested primary’ and ‘non-forested primary’; and natural land – the sum of ‘potentially forested secondary’, ‘potentially non-forested secondary’, ‘managed pasture’ and ‘rangeland’. To estimate Primary habitat loss we found the mean primary habitat across sampling points in each population area for each year in the population monitoring period. We then estimated the rate of loss in primary habitat over time by dividing the rate of loss in each year by the previous year, and then converted this to a percentage loss. We defined the mean Primary habitat loss (%) for each population area as the average across this timeseries of loss rates. We followed an identical procedure for Change in natural land. Importantly, primary habitat cannot be restored in the time spans we considered, so our estimate of change in primary habitat could only decrease or remain stable. Whilst natural habitat can fluctuate up and down.

We estimated the Change in human density using the Global human settlement human population raster ^4^, which describes the human density per km^2^ for four years: 1975, 1990, 2000, 2015. For each year, we reduced the spatial resolution to 0.1° by averaging (mean) over finer-resolution pixels. In order to estimate the Change in human density for each population area’s monitoring period, we had to estimate missing human density values (years) in each population area. To do this, we first extracted the mean human density across each population area, in all four of the available years. In each population area, we then used a log-linear regression to model human density (base log transformed) against year, and then predicted human density between 1960 and 2015. As human density change was non-linear we modelled year (predictor) with a cubic fit. We then extracted the back-transformed predicted values of human density for all years in each population area. As we were only working with four data points, model predictions were highly uncertain. This uncertainty was included by resampling our model with 100 bootstrap iterations. For each population monitoring period and iteration, we extracted the predicted human densities and estimated the rate of change (%) as calculated for the other land-use covariates. Finally, we calculated the mean human density rate of change (%) across all iterations, as well as the standard deviation, which was used to represent uncertainty in the values within the inference model (see below).

##### Climate

Our two climatic covariates, Change in extreme heat and Change in drought, describe how the number of months exceeding an extreme heat or drought threshold (respectively) changed between a pre-industrial period (1850 – 1900) and the population monitoring period (Figure S2). To derive these Change in extreme covariates, we compiled a raster timeseries of bias-corrected daily maximum near-surface air temperature from 1850 to 2014, at a 0.5° resolution ^5–7^. We then averaged the daily maximum temperature for each month, in each year, creating a monthly raster timeseries of the mean maximum temperature from 1850 to 2015. From this, we calculated the mean and standard deviation of the monthly maximum temperature in the pre-industrial period, and defined the extreme heat threshold as the mean plus 2 standard deviations of the mean in each pixel. With this approach, approximately 2.5% of the monthly values in the pre-industrial period would exceed this threshold, thus, representing rare extreme heat events. Next, we quantified the actual mean number of months per year in the pre-industrial period that exceeded this threshold, as well as the number of months to exceed this threshold in all years between 1960 and 2015 for every pixel. We then subtracted the number of threshold-exceeding months in each year (1960-2015) from the mean threshold-exceeding months in the pre-industrial period, creating a raster timeseries describing the difference in threshold-exceeding months. This value could inform us about how the number of months exceeding the threshold in a given year in which a carnivore population was studied differ to the average across the pre-industrial period. Finally, for each sampling point in each population area, we found the mean difference across the population monitoring period, and then averaged this difference across all sampling points to produce a population area estimate of the Change in extreme heat.

To derive our Change in drought covariate, we compiled two raster timeseries’ describing the bias-corrected daily near-surface air temperature and daily bias-corrected precipitation, both from 1850 to 2014, at a 0.5° spatial resolution ^5–7^. We then averaged the daily temperature rasters and summed the daily precipitation rasters for each month, in each year, creating two monthly raster timeseries, describing monthly mean temperatures and total precipitation (in mm) from 1850 to 2015. For temperature, we calculated Thornthwaite’s evapotranspiration across the raster timeseries, which uses the mean temperature, latitudinal position and number of daylight hours to estimate the evapotranspiration rate ^8^. Next, we subtracted this monthly evapotranspiration estimate from the monthly precipitation (mm) estimate to produce Thornthwaite’s standardised precipitation-evapotranspiration index (spei), a standard metric used to describe water availability ^9^. We then proceeded to estimate a spei threshold and the mean difference in months overlapping the threshold (pre-industrial vs. population monitoring period) in an identical way to how monthly maximum temperature was used to estimate the Change in extreme heat covariate.

##### Governance

We identified five governance covariates that we considered important to large predator population trends, four of which were measured at the country-level: War present, National governance, Human development, and Change in human development. We used three datasets to populate these covariates: 1) For War-present, we used the UCDP/PRIO Armed conflict dataset ^10^, which lists conflicts (between 1946-2019) where fatalities per year exceeded 25 and at least one of the parties was governmental. We summarised this dataset into a timeseries that describes whether a war was taking place in each country’s territory in each year between 1960 and 2016. 2) For Governance, we extracted the world governance indicator metrics ^11^, which present six annual governance timeseries for each country between 1996 and 2016. 3) Finally, for Human development we sourced the UN human development index ^12^, which provides an annual timeseries describing life expectancy, education level, and income per capita between 1990 and 2016 for 189 countries.

As the governance and human development indicator data only stretch back until 1996 and 1990, respectively, some of the trend data preceded the indicator values. We imputed missing values through a multiple imputation chained equations (MICE) framework ^13^. We used a hierarchical normal (2l.pan function) imputation model, where observations are nested into the different countries, and included: the year of the observation, the six governance indicator metrics, the human development index, whether war was present in that year (yes or no), as well as the country’s gross domestic product (log 10 transformed). MICE imputations are stochastic and repeated numerous times, creating an approximate distribution for each missing value. We imputed missing values for each variable between 1960 and 2016 in each country, and repeated the imputation 100 times with a 50 iteration burn in – all variables showed convergence (Figure S4).

**Figure S4.**
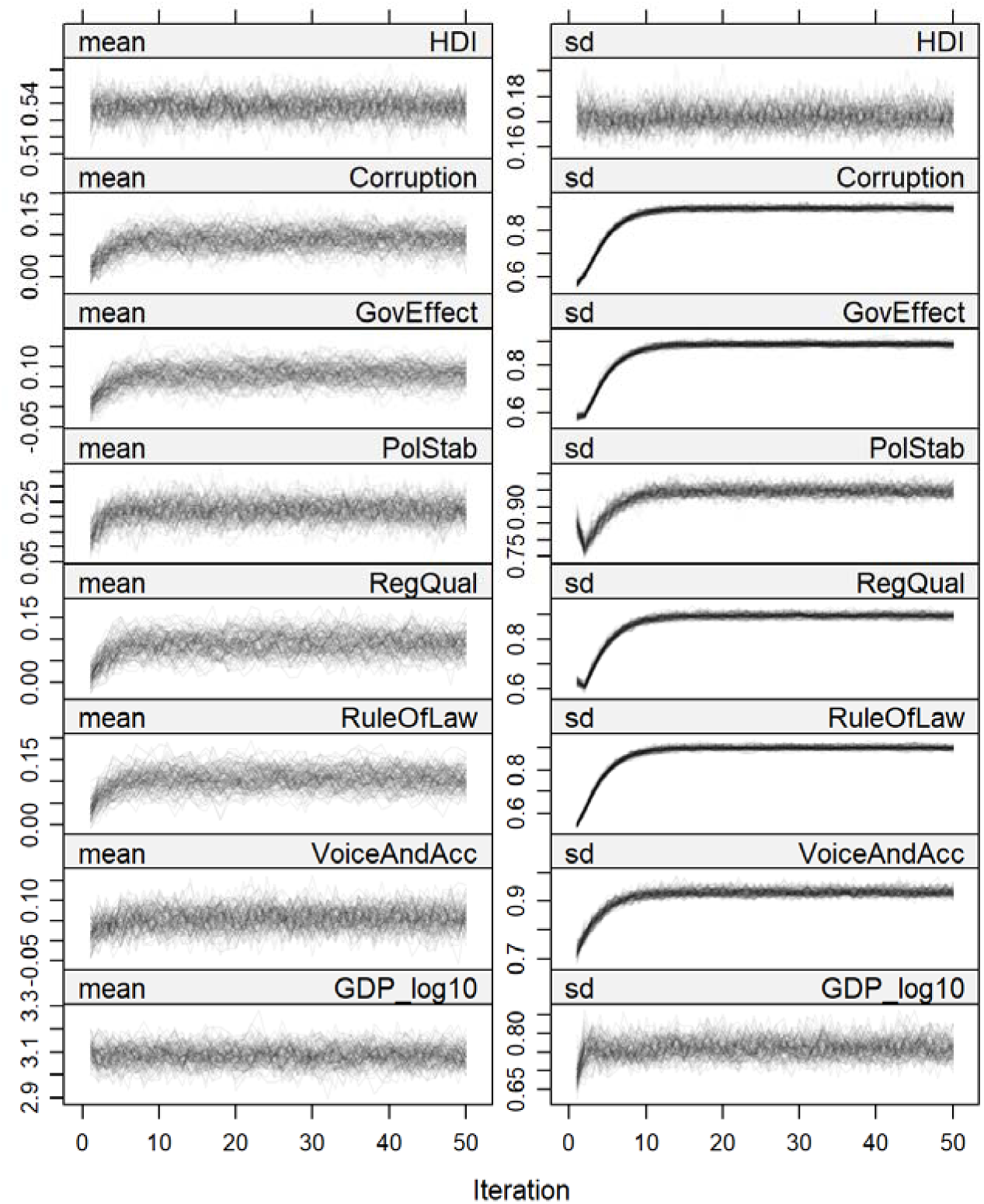
Convergence of mean (left) and standard deviation (right) of variables with missing values in the imputation model: HDI – human development index, Corruption – control of corruption, GovEffect – government effectiveness, PolStab – political stability and absence of violence, RegQual – regulatory quality, RuleOfLaw – rule of law, VoiceAndAcc – voice and accountability, and GDP_log10 – gross domestic product (log 10 transformed). Convergence ran with 50 iterations and 100 chains.

Using the imputed datasets, we extracted the mean value across the six governance indicators in each country, year, and imputation chain. We then calculated the mean and standard deviation of this combined governance across the imputation chains to produce an annual governance timeseries (and associated error) for each country. For Human development, we averaged over the 50 stored imputation chains to calculate the mean and associated standard deviation for each country and year. We assessed if imputed values were plausible (Figure S5).

**Figure S5.**
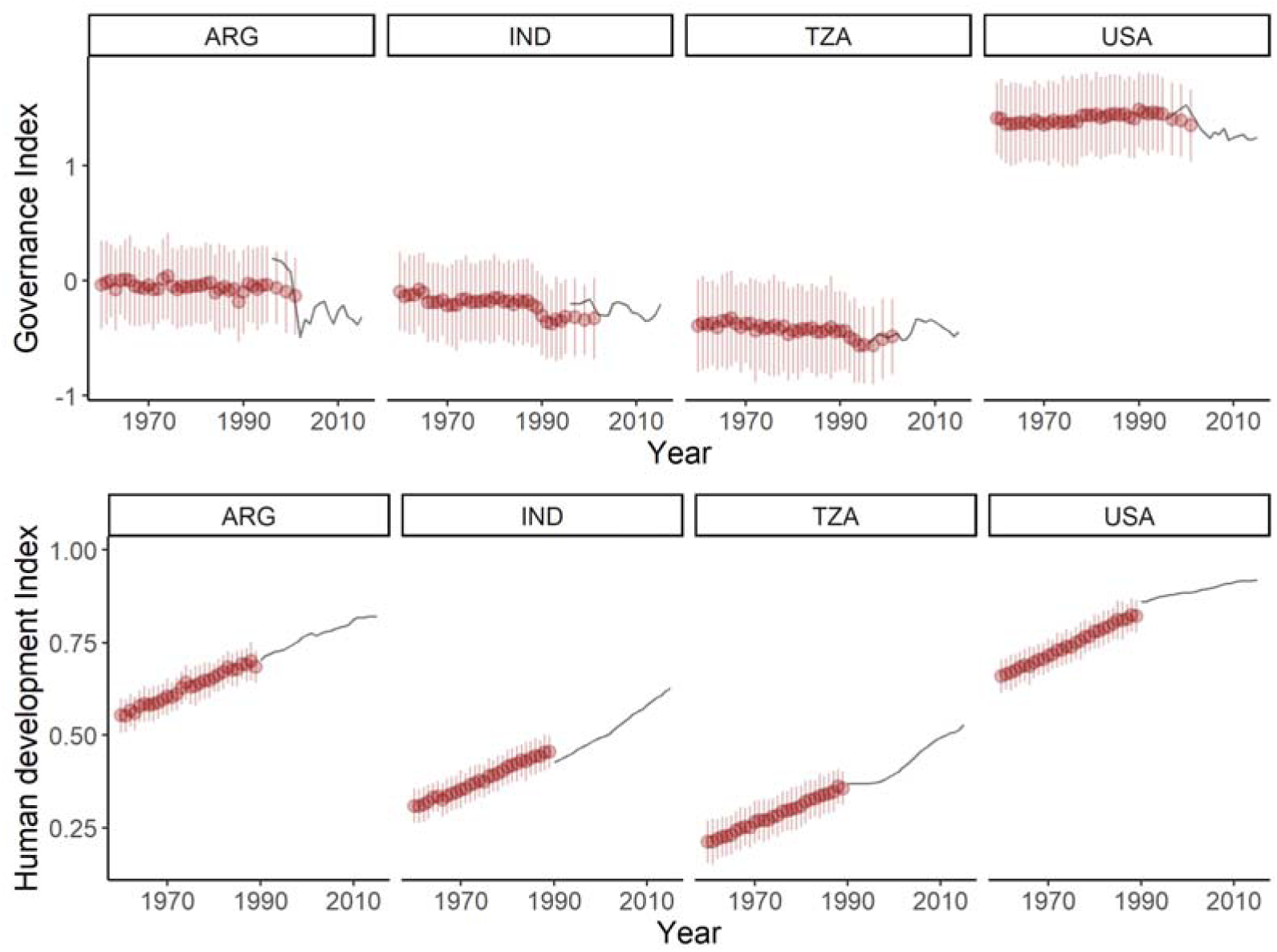
Governance (top) and human development (bottom) index scores for Argentina (ARG), India (IND), Tanzania (TZA), and the United States of America (USA). True values are depicted with the black line, whilst the mean imputed values (point) and associated 95% confidence intervals (bars) are depicted in red.

After we derived the governance and human development timeseries, we began extracting the covariates. For War present, we created a binary variable that described whether war(s) had occurred in the country where the population is located, at any point during the population monitoring period. For Governance and Human development, we extracted the mean scores per country, and associated standard deviations from the final year of the population monitoring period. For Change in human development, we extracted all human development values across the population monitoring period, and divided each value by the value in the previous year to produce a timeseries describing the annual changes in human development. We then averaged these values and converted the average into a percentage which describes the annual rate of change (%) in human development.

Our only governance covariate not measured at the country-scale is Protected area coverage. For this variable, we compiled the annual timeseries of protected areas polygons covering the period 1960 to 2020 from the World Database of Protected Areas ^14^. In each year, we converted the polygons into a 0.1° resolution raster describing the proportional cover of protected areas in each pixel. In the final year of the population monitoring period, we calculated the mean coverage of protected areas across pixels within the population area.

##### Traits

We identified five species traits which could influence population trends in large predators: Body mass, Maximum longevity, Climatic niche breadth, Ecological niche breadth, and Reproductive output. Body mass describes the average body weight of an adult of the species in grams (log_10_ transformed), Maximum longevity describes the maximum lifespan of an individual of the species in years (log 10 transformed), and Climatic niche breadth describes the standard deviation of the mean monthly temperatures across the species current IUCN range, calculated using WorldClim 2.1 ^15^. The other two trait-covariates, Ecological niche breadth and Reproductive output, are principal components of a larger array of traits. Specifically, Ecological niche breadth captures habitat and diet breadth. Habitat breadth is defined as the frequency of different IUCN habitat classifications ^16^ the species occurs in. Diet breadth is defined as the number of different food-types the species has been recorded consuming (or with evidence of consuming through faecal or stomach content analysis), from the following 12 options: mammals, birds, reptiles and amphibians, fish, invertebrates, fruit, pollen and nectar, leaves and branches, seeds, grass, roost and tubers, and carrion – sourced from an unpublished trait dataset ^17^. Our Reproductive output trait is a principal component of the following traits (all log 10 transformed): interbirth interval, gestation length, litter size, minimum breeding age, neonatal body mass, and weaning age. As a result, our five traits of interest were reliant on collecting values for 12 common traits.

We sourced values for our traits from three different trait datasets: PanTHERIA ^18^ AnAge ^19^, and an unpublished large predator trait dataset ^17^. We used multiple trait datasets to populate missing values at the species level. However, the values sometimes differed between the trait datasets, and in these cases, we created multiple records for the species to capture this uncertainty in the trait value. As a result, many species had more than one value for a given trait. However, despite using multiple trait datasets, values were still missing for some species in some traits (Table S1), and so we imputed missing trait values with Rphylopars ^20^. Rphylopars outperforms MICE imputation (used in the governance covariates above) when imputing species traits as it uses both the trait values and species’ phylogeny to estimate missing values – Rphylopars is considered one of the best imputation methods ^21^. In our Rphylopars model, we trialled three ^22^ Carnivora phylogenies to ensure the imputations did not drastically change depending on the phylogeny (Figure S6). Once we confirmed the choice of phylogeny had little impact, we proceeded with the rest of the analysis only using the phylogeny considered ‘best’ by ^22^. We also included all 12 traits mentioned above in the imputation model, and six other traits to attempt to account for biases in the imputation model, specifically: species area of occurrence, minimum absolute latitude species occurs at, maximum absolute latitude species occurs at, difference in maximum and minimum latitude, maximum mean monthly temperature species occurs at, and minimum mean monthly temperature species occurs at. As the phylogenies we used were not perfectly matched to the CaPTrends and Living Planet Index taxonomies, we corrected synonymous species names in the phylogeny, and where species included in the taxonomy were absent from the phylogeny, we appended the species to a close relative node (inferred from taxonomy).

**Table S1.** Percentage of values missing in each trait. The six additional traits added simply to minimise bias had no missing values and so are excluded from this Table.

| Trait | Missing trait values (%) |
| --- | --- |
| Body mass | 8.0 |
| Maximum longevity | 12.6 |
| Climatic niche breadth | 0.0 |
| Habitat breadth (part of ecological niche breadth) | 0.0 |
| Diet breadth (part of ecological niche breadth) | 23.0 |
| Age of sexual maturity (part of reproductive output) | 24.1 |
| Litter size (part of reproductive output) | 10.3 |
| Gestation length (part of reproductive output) | 9.2 |
| Weaning age (part of reproductive output) | 27.6 |
| Interbirth interval (part of reproductive output) | 12.6 |
| Neonatal body mass (part of reproductive output) | 28.7 |

**Figure S6.**
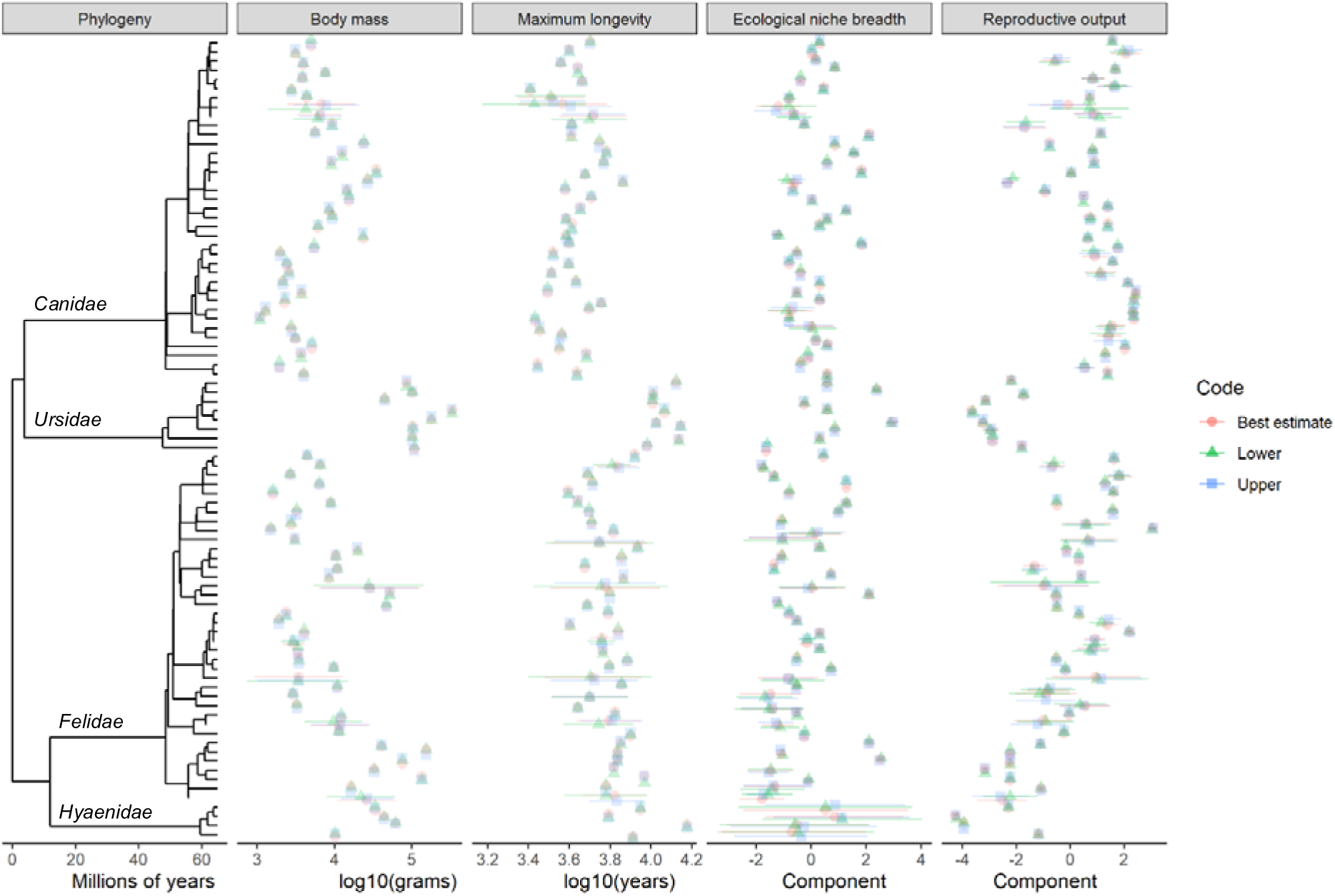
Trait values represented on the phylogeny; climatic niche breadth is excluded as it had no missing values. Error bars represent the 95% confidence around the mean imputed values. If observations were complete (i.e. not missing values) the standard deviation around the observation was zero and so there are no confidence intervals. We include the three phylogeny types in ^22^.

An advantage to using Rphylopars is that it provides an estimate of the standard deviation around the missing trait values, which is something we wanted to capture in our modelling (see Inference model below). Three of our traits were used as covariates directly within the modelling (Body mass, Maximum longevity, and Climatic niche breadth), so required no further manipulation as their associated standard deviations were available from the imputation. However, Ecological niche breadth and Reproductive output required dimension reduction through principal component analysis (PCA), with the number of variables shifting from 2 to 1, and 6 to 1, respectively. Performing PCA on the mean values would fail to capture trait uncertainty, and so instead we developed normal distributions for each species’ trait value using their mean, and an error of one standard deviation. We then sampled from each distribution 100 times, and each time conducted a PCA on the trait to develop an eigenvector. We saved the eigenvector values on each of the 100 repeats, and once the repeats were complete, we calculated the mean and standard deviation for each species across the eigenvector values. This PCA sampling procedure was performed separately on the Ecological niche breadth and Reproductive output components. We examined the trait values to ensure they were plausible (Figure S6), and also checked between-trait correlations were acceptable i.e. sufficient variance in the correlation (Figure S7).

**Figure S7.**
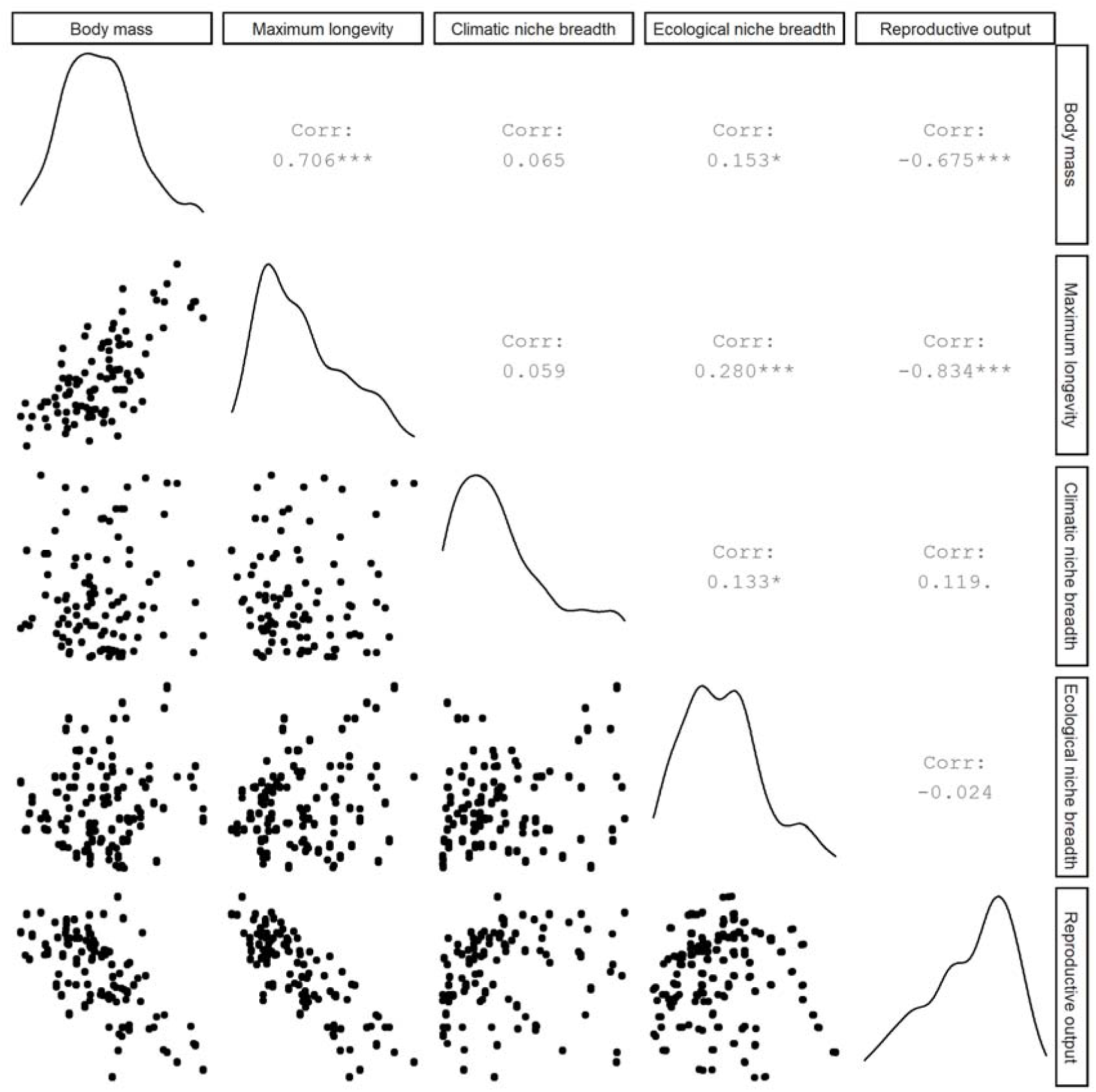
Distributions of traits are represented on the diagonal. The person correlations between traits are represented above the diagonal with varying levels of statistical significance (p-value): ‘***’ when p<0.001, ‘**’ when p<0.01, ‘*’ when p<0.05, and ‘.’ when p<0.10. The scatter of observations (one point per species) below the diagonal graphically represents these relations.

##### Temporal lag

A challenge in identifying how covariates impact population trends, is matching the temporal scale of the covariate with the population i.e. How long is the lag between the covariate impact and a change in the population?. This lag period is likely variable across covariates (i.e. it could be different with land-use and climate features) and species traits. For example, recent work has suggested population change in large mammals peaks at approximately 8 years after forest loss ^23^. As a result, we conduct sensitivity analysis (see Sensitivity analysis below) to determine how model fit was influenced by lag selection, considering three options: 1) No lag, so covariate changes are measured between the start and end year of each population monitoring period. 2) Five-year lag, where covariate changes are measured between the five-years prior to the start of each population monitoring period, and run to the end of each period. 3) Ten-year lag, where covariate changes are measured between the ten-years prior to the start of each population monitoring period, and run to the end of each period.

##### Cleaning data

We opted to remove a selection of the population trend and covariate data as the values were deemed unreliable or unsuitable. Specifically, we removed any population trend records beginning before 1970 or after 2016 (N = 11), where governance data was largely incomplete. We also removed records overlapping multiple countries (N = 10), and any population trends with an excessively large population buffer-area (N = 40) – we set the threshold at 2 million km^2^ which could accommodate state and small-country level estimates, but would exclude large countries. For example, the largest population area in the dataset covered all of Russia (∼21 million km^2^). Any population trends discussing non-native species were removed (N = 6), as well as records not overlapping any land (N = 4) e.g. *Ursus maritimus* populations occurring exclusively on sea-ice. We also removed any population trends where the population had either recolonised an area or become locally extinct (N = 80), which represent an extreme form of population change that could skew our inference. After excluding records, we were left with 989 estimates of annual rate of change, and 138 qualitative descriptions of change.

#### Modelling - Inference model

We fitted a hierarchical linear model (Figure S8) to determine the effect of a combined 23 covariates and interactive effects on the rate of change in large predator populations. Our model development falls into seven compartments: response, random intercepts, coefficients and covariates, imputation uncertainty, weighted error, confirming parameters, and model running. The model was written in BUGS language and implemented in JAGS 4.3.0 ^24^ via R 4.0.3 ^25^.

##### Response

The core of our model is a linear regression with fixed and random effects attempting to predict some latent state (unknown annual rates of change; Equation 1). Our observed annual rates of change are realisations of this latent state and are linked to linear model predictions through a state-space structure (Equation 2).

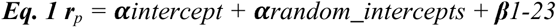

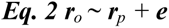

Where ***r****_p_* represents the deterministically predicted latent annual rates of change from a linear model (simplified here) containing an intercept and random intercepts (α), as well as fixed effect beta (β) coefficients. ***r****_o_* represents the observed (or realised) annual rates of change which are stochastically linked to a normal distribution of ***r****_p_*. The uncertainty in the normal distribution is determined by error e. When building the model, we identified that the model residuals exhibited a heavy tailed t-distribution and so we transformed our responses into a gaussian distribution with an inverse-hyperbolic sine transformation.

**Figure S8.**
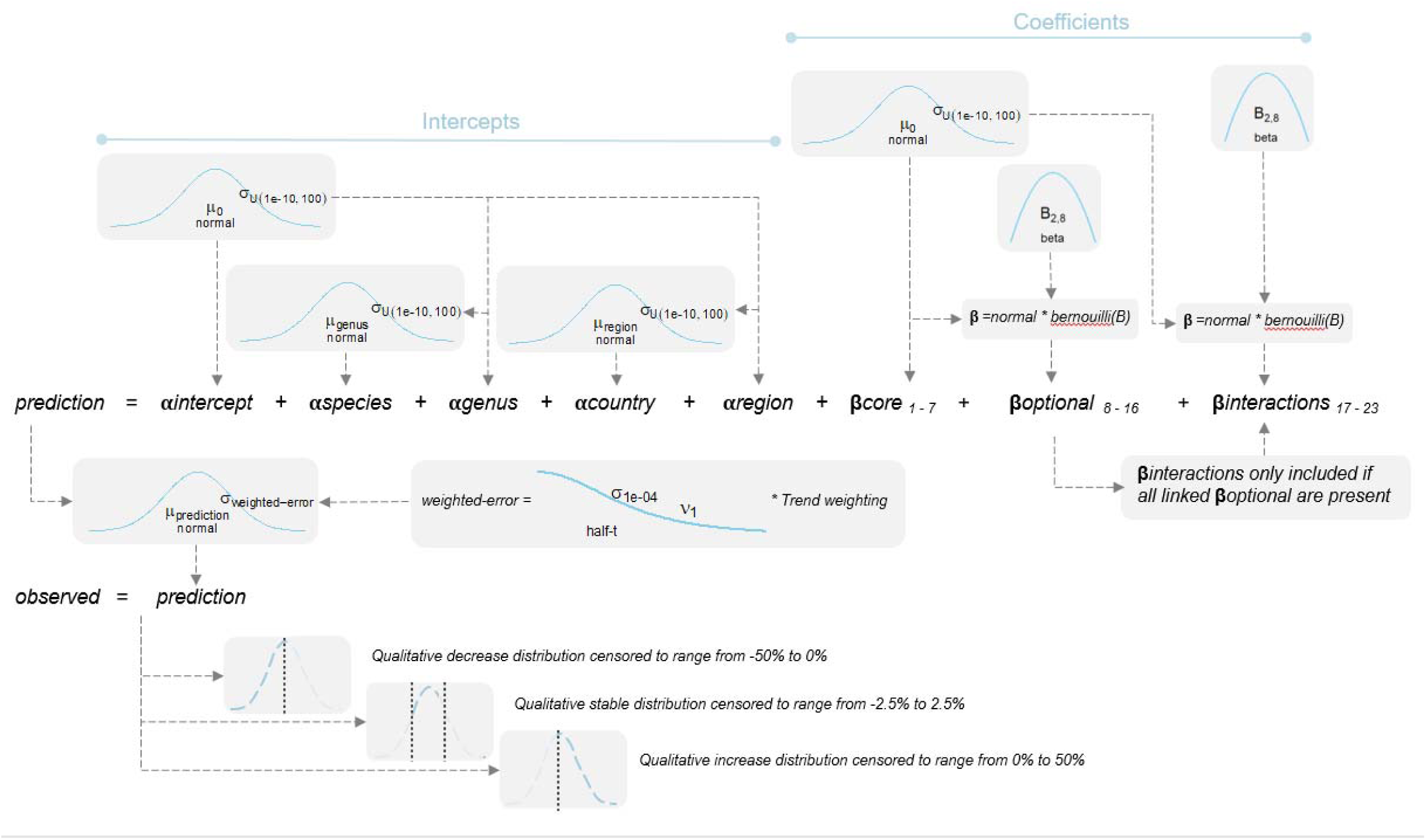
Model structure of hierarchical linear model, describing distributions of priors and hyperpriors, as well as the process for incorporating overall error, imputation error, trend weights, and censoring within the model. We use five distributions (parameters described in brackets) within the model: normal (μ = mean, σ = standard deviation), beta (shape1, shape 2), half-t distribution (μ = mean, σ = standard deviation, df = degrees of freedom), U/uniform (minimum, maximum), and Bernoulli (B = probability).

Our observed population trends fall into two types: quantitative annual rates of change and qualitative descriptions of change. Both data types were modelled with the same normal error prior (Eq2), but to deal with the different data types, and the unknown values of the qualitative descriptions, we censored the qualitative records to indicate that the true value is unknown, but it occurs within a specified range. We specified these annual rate change ranges as -50% to 0%, -5 to 5%, and 0% to 50% within the decrease, stable and increase categories, respectively.

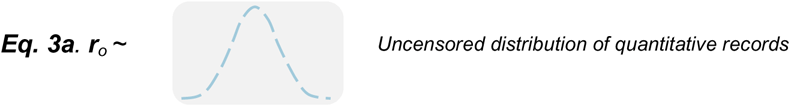

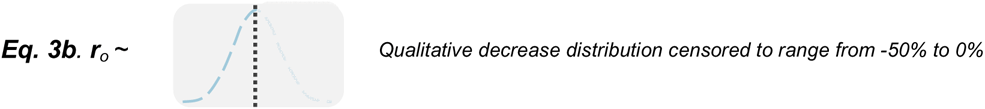

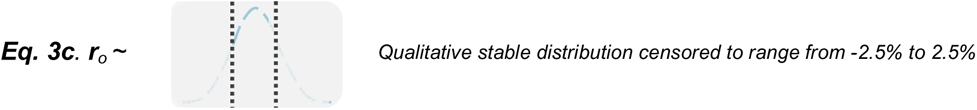

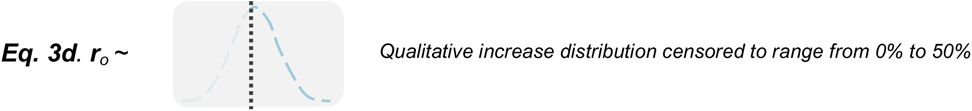

Equation 3 is a distributional representation of Equation 2 describing how the observed annual rate of change is constrained (or censored) to occur within specific values.

The censoring range thresholds are similar to the range of the observed rates of change (−75% to 68%). Many of the qualitative records occur in less-well represented regions, species, and time-periods, so their inclusion addresses known data biases (Figure S1). However, these lower quality records can be more prone to error. As a result, we conduct sensitivity analysis (see Sensitivity analysis below) to assess how including censored observations altered model fit, compared to only using quantitative, and high-quality quantitative (derived from at least three abundance observations), trends.

To allow the latent rates of change to vary (***e*** in Equation 2), we set the standard deviation of the latent states normal distribution as a half-t (or half-cauchy) hyperprior (centred at zero, with a standard deviation of 0.001 and one degree of freedom). However, as some observations were likely to be more robust than others, we altered the standard deviation of the latent state normal distributions depending on each observation’s quality. We varied the standard deviation by multiplying the half-t hyperprior by a deterministic weighting term, where the standard deviation was inflated for lower quality observations (see *Weighted error* below).

##### Random intercepts

We used a hierarchical model structure to account for phylogenetic and spatial non-independence in the data, including species as a random intercept nested with genus, and country as a random intercept nested within sub-regions, as defined by the United Nations (https://www.un.org/about-us/member-states). These parameters were fit with a normal distribution centred at zero and their error terms were given a vague uniform hyper prior, with a standard deviation ranging from 1*e*^-10^ to 100.

##### Coefficients and covariates

With a combined 23 covariates and interactive effects, we were conscious of overparameterizing the model. As a result, we split these parameters into three groups: 1) core parameters – which included main effects that have previously been reported as influential, are expected to be influential, or control for other parameters and methodological features; 2) optional parameters – which included main effects we considered interesting but with little evidence to-date of any influence on trends; and 3) interaction parameters – which includes all interaction terms between parameters. Core parameters included: Change in human density, Primary land loss, Population area, Body mass, Change in extreme heat, Governance, and Protected area coverage. These core parameters were included in every model, but we used Kuo and Mallick variable selection ^26^ to identify important parameters from the optional and interaction groups, where variables were only included in an iteration if they were selected from Bernoulli priors. Our optional parameter group was assigned a Bernoulli prior, which sampled from a beta hyperprior (*a* = 2, *β* = 8), such that approximately 20% of optional effects would be included in any iteration, on average, but this could range from 0 – 100%. The interaction parameter group had an identical, but separate prior setup. Crucially, this interaction prior was only activated if both main effect parameters were present in the model. For example, for the Change in extreme heat and Change in drought interaction to be selected, it would require Change in drought to be selected from the optional Bernoulli prior, and then the interaction itself would need to be selected from the interactive Bernoulli prior. As variable selection can be highly influenced by the standard deviation of the parameter slope coefficients, we specified the slope standard deviation as a vague uniform hyperprior ranging from 1*e*^-10^ to 100.

##### Imputation uncertainty

Six of the covariates in the model contained missing values that were filled using imputation (see *Land-use*, *Traits* and *Governance* within the *Covariates* section above). To improve the robustness of our model inference, we accounted for uncertainty in the imputed estimates by treating imputed values of the covariates as distributions instead of point estimates, where each imputed value was assigned a normal distribution centred at the mean imputed estimate and with an error varying by the imputed observation standard deviation. As we z-transformed all of our covariates to standardise coefficients, except ‘War present’ which is a categorical variable, we also had to rescale the associated imputation standard deviations. As standard deviations cannot be rescaled in the same way as the imputed estimates, we first converted the standard deviation into confidence intervals, we then z-transformed the intervals using the mean and standard deviation of the covariate, and then back calculated the standard deviation from these intervals.

##### Weighted error

When developing the model, we were conscious that all not rates of change should contribute equally to the fit. For example, whilst including the censored records could decrease taxonomic and spatial biases in the data, they may also introduce error, as these censored records are unlikely to be as accurate as the quantitative trends. As a result, we included a weight term to inflate the uncertainty in these lower quality records, where the half-t hyperprior discussed above is multiplied by a weight term defined as the inverse of the estimated error in the rate of change. This weight term was developed through simulation (see below), and these simulated error weights inflated the variance around the trend in all low-quality observations, not just the qualitative ones.

When simulating the trend weights, we considered our real trend data to be estimates of true trends with some degree of error. This error would be influenced by the certainty of the population abundance estimates, the sampling intensity (e.g. Is the population sampled every year or only in 50% of years?), and the sampling duration (e.g. Is the trend based on 2 or 20 years?). As a result, we developed a simulated trend dataset comprised of ‘true’ trends where abundance values are known (not estimates) and complete, and an edited trend dataset where abundances are uncertain and missing as expected in a real scenario and observed in our trend dataset.

For the ‘true’ trend dataset, we simulated 6000 timeseries of abundances which varied in duration (from 2 to 20 years), with an estimate of abundance in all years throughout that duration. We then calculated the true trend for each timeseries by modelling abundances (response) against year in a log-linear regression, and converted the slope estimate into an annual rate of change (%). Abundance values exhibit a normal distribution ranging from approximately 0 to 500.

For the edited trend dataset, we altered two parameters in each of the 6000 timeseries of abundances generated above. Firstly, for each abundance estimate in each timeseries, we developed a random normal distribution, centred on the true abundance value, but with varying levels of error (coefficient of variation from 0.02 to 0.2). For example, with a true abundance of 100, a low error of 0.02 would produce a range of abundance estimates from approximately c.95 to c.105, whilst the abundance would range from c.50 to c.150 with an error of 0.2 (Figure S9). We sampled from these newly created abundance distributions to produce new error-prone abundance estimates, reminiscent of real uncertainty in abundance estimation. Secondly, we removed a random sample (between 0% and 90%) of the observations in each timeseries, producing timeseries’ with varying levels of completeness. We then re-calculated the annual rate of change (%) as in the true trend to produce the estimated trend. The distribution of the estimated trend was largely similar to the true trend obtained from the complete dataset (Figure S10).

**Figure S9.**
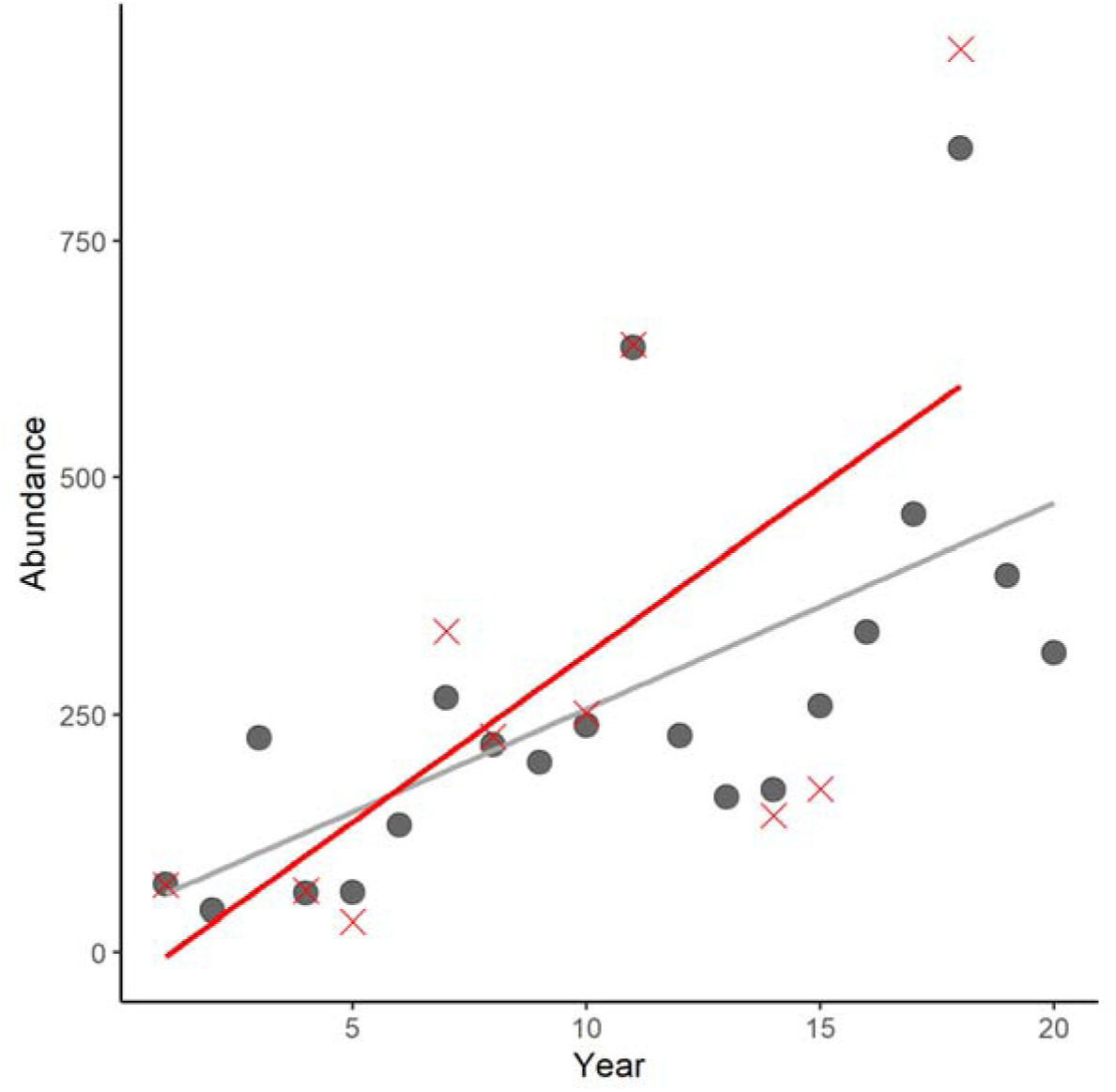
Impact of adding noise to abundance values and removing abundance values on the population trend, with the true trend (derived from known and complete abundance values) in grey, and the estimated ones in red. In this example, the coefficient of variation equals 0.2, with a sampling intensity of 50% i.e. half the years in the population monitoring period have abundance values.

**Figure S10.**
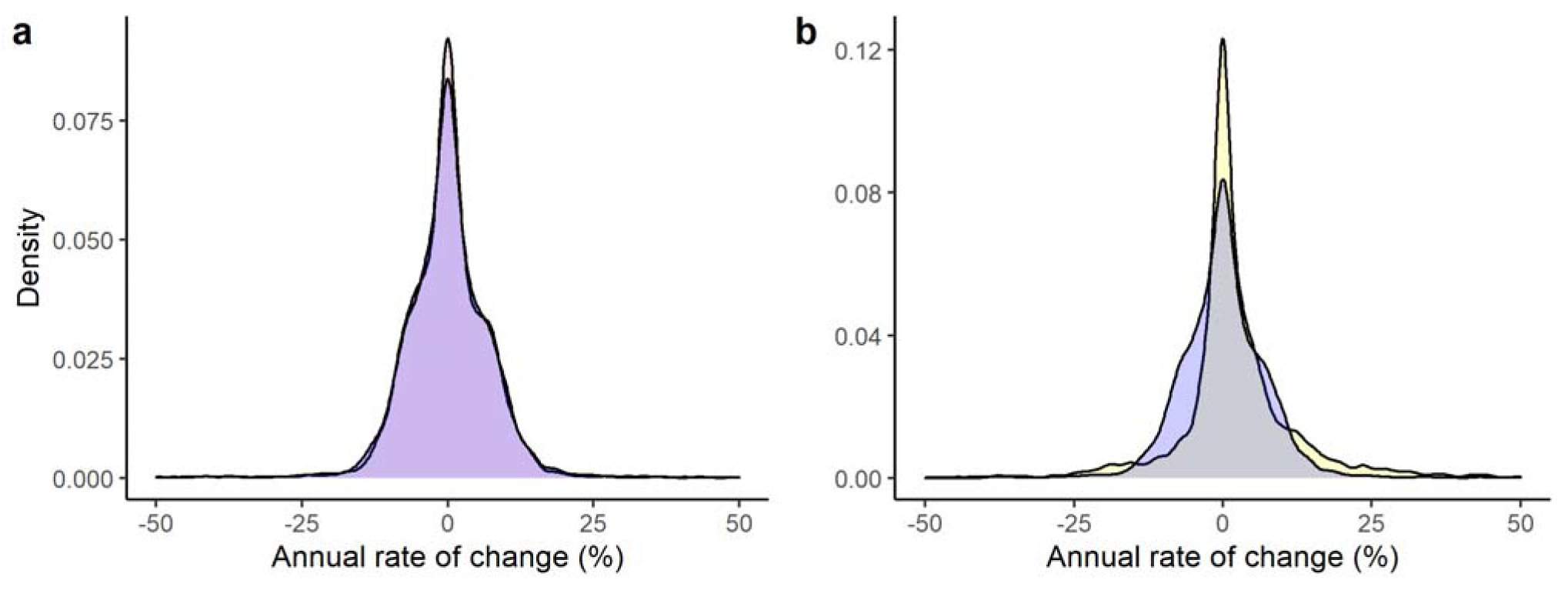
a) Distribution of simulated true trend values (pink) and simulated estimated trend values (blue). b) Distribution of simulated estimated trend values (blue) and real trend values compiled from CaPTrends ^1^ and the Living Planet Index ^2^ in yellow.

We extracted the absolute error (difference) in the annual rate of change of the true and estimated trends, and modelled this error (as the response) against sampling intensity (what percentage of years have observations), the coefficient of variation around the abundance estimates, and the duration of the trend, all in a log-linear regression. Trends with a higher sampling intensity (coef = -1.59, CI: -1.72, -1.47), lower coefficient of variation (coef = 4.09, CI: 3.68, 4.49), and longer duration of the trend (coef = -0.12, CI: -0.13, -0.11), had lower errors (Figure S11). We used this model based on simulated data to predict the likely error in the real data. For sampling intensity, we calculated the percentage of abundance values used to calculate the trend relative to the trend duration. For the trend duration, we calculated the number of years in each population monitoring period. Unfortunately, in most cases the estimates of uncertainty around the raw abundance values were unavailable, so we were unable to directly calculate the coefficient of variation for each trend. However, we did have data describing the quality of the sampling and modelling which could act as a proxy for the accuracy of the abundance values. Specifically, we scored trends separately in three areas (Table S2), where trends could only be assigned to one category per area; we then added the score across the three areas: *Sampling* – how systematically was the population sampled? *Modelling* – how robust was the approach for modelling abundance values? *Low-quality record* – does the record meet any of the criteria for being considered low quality? For example, a trend with systematic population sampling (+0.04), in which sampling effort was accounted for (+0.04), and none of the low-quality criteria were met (+0), would be given a coefficient of variation score of 0.08. For an abundance value of 100, this coefficient of variation score would allow the abundance to vary between 75 and 125. Admittedly, our scoring criteria are arbitrary, simply designed to add uncertainty around trends that used less robust methods, rather than describe the true uncertainty in the trend. However, as these arbitrary values only contribute one feature of three in the weighting system, there impact is likely minimal, and is tested in sensitivity analysis regardless (see below).

**Figure S11.**
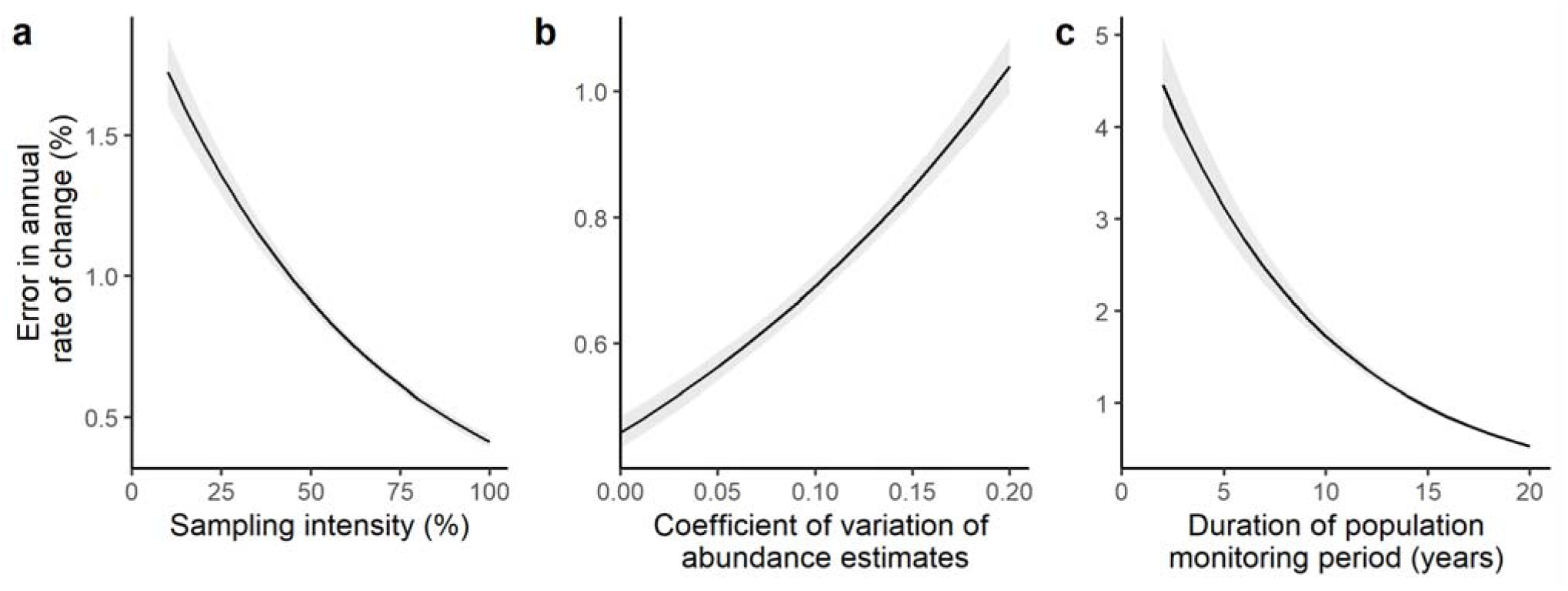
Marginal effect of sampling intensity (a), coefficient of variation around abundance values (b), and trend duration (c) on the absolute error in the annual rate of change (%), comparing the simulated-true to the estimated trend. Sampling intensity describes the percentage of years with abundance values in the population monitoring period. Trend duration describes the length of the population monitoring period e.g. 1990 – 1992 equals three years.

**Table S2.** Scoring criteria used to define a coefficient of variation (CV), uncertainty, in abundances.

| Description |  | CV |
| --- | --- | --- |
| *Sampling* | Method of population sampling is not described or is unsystematic/biased. | 0.08 |
|  | Method of population sampling is systematic. | 0.04 |
|  | All individuals in the population identified. | 0.01 |
| *Modelling* | Method of deriving abundance from population sampling is not described or values are just reported in their raw format. | 0.08 |
|  | Sampling effort accounted for in abundance estimates. | 0.04 |
|  | Abundance derived through complex modelling, or total abundance known. | 0.01 |
| *Low quality record* |  |  |
|  | Abundance values derived from genetic or harvest data; or the trend is labelled as inaccurate within the primary literature; or trend describes asymptotic instead of observed growth; or trend metric is unconventional. | 0.04 |

After predicting the error in the real trend data using the simulated weight model, we scaled and flipped the values so that 1 indicates low error and 1e-10 indicates high error. These values had to be flipped, as the weight term in our hierarchical linear model (Figure S8) is multiplied by the precision (i.e. uncertainty) around each trend observations, in which a precision would then be deflated (i.e. uncertainty inflated) if multiplied by a high error trend. For example, for observation A with a low error of 0.9, a precision of 10 would be deflated to 6, whilst for observation B with a high error of 0.1, a precision would be deflated to 1, so A would receive 6 times more weight than B.

To ensure our weight term benefitted the model fit, we conducted sensitivity analysis to compare the model fit under four options: 1) the simulated error weight (described above); 2) weighting by trend sample size, whereby trends derived from more abundance observation are given more weight; and 3) unweighted i.e. all observations are treated equally.

##### Sensitivity analysis

We conducted sensitivity analysis to test how the different weighting, censoring, and temporal lag options influenced our model results, with the aim of selecting parameters which maximised model marginal and conditional R^2^, whilst also balancing this decision against potential risks. For example, including censored observations may reduce model fit but this could still be worthwhile if it reduces taxonomic and spatial biases. For weighting, we ran models separately under each of the three options, including censored observations and a 5-year lag on all covariates in all cases. After identifying the simulated error weighting as the best option for maximising fit and minimising bias (see Supplementary results) we tested the censoring options, again holding all covariates at the 5-year lag. Including censored observations was valuable, so we included the censored observations when assessing the different temporal lag models, from which we identified that using a 10-year lag improved model fit. In each case, we ran the model through two chains, each with 10,000 iterations and discarding the first 5,000. We thinned the complete chains to store every other iteration (thinning factor of 2). We monitored convergence of key parameters within each model, specifically: standard deviation of the model intercept, standard deviation of beta coefficients, standard deviation of each random effect (regions, countries, genus and species), standard deviation of the overall model error, the optional parameter beta hyperprior, and the interactive parameter beta hyperprior. We ensured the multivariate potential scale reduction factor was less 1.1 across all models in the sensitivity analysis.

##### Model running

After selecting the simulated error weighting, censored observations, and a 10-year lag from the from the sensitivity analysis (see Sensitivity analysis in the supplementary results), we ran the full model through three chains, each with 120,000 iterations. The first 20,000 iterations in each chain were discarded, and we only stored every 10^th^ iteration along the chain (thinning factor of 10). We opted for a large chain and burn-in due to the model complexity, and to allow a broad selection of parameter combinations to be tested under variable selection. We assessed convergence of the full model on all parameters monitored in the sensitivity analysis, as well as the model intercept, and all 23 main and interactive effect slope coefficients. We checked the standard assumptions of a mixed effect linear model (normal residuals and heterogeneity of variance), and tested the residuals to ensure no spatial (Moran’s test) or phylogenetic (Pagel’s lambda) autocorrelation. We also conducted posterior predictive checks to ensure independently simulated values were broadly reminiscent of model predicted values.

After model running, we calculated how frequently, as a proportion, each of the 23 main and interactive effects occurred within the iterations. For the optional parameters, this was derived by dividing the frequency of occurrence by the total count of iterations. For the interactive parameters, whose inclusion was dependent on the frequency by which their derivative main effects were selected, we divided the frequency of occurrence by the total count of iterations where both derivative main effects were present. We also report the median slope coefficient and associated credible intervals for each of the main and interactive effects, and produce marginal effect plots for a selection of important parameters. These marginal effects hold all other covariates at zero (which is the equivalent of the mean, as covariates were z-transformed). We also display the distribution of the random intercepts e.g. for each region, country, genus and species.

### Results

#### Sensitivity analysis

We assessed how trend weighting, including censored observations, and specifying a lag period on the covariates influenced model fit and inference, in part to assess if results were particularly sensitive to specific parameters, but also to help choose the parameters which optimised fit and spatio-taxonomic coverage. Using censored observations and a lag period of 5 years on covariates, model fit was greater when using the simulated error weights compared to the unweighted model and the model weighted by sample size (Table S3). Using simulated error weights and a lag period of 5 years on covariates, we then tested the impact of including censored observations which showed higher marginal and conditional R^2^ when censored observations were included. Using only high quality timeseries (compared to including censored observation) resulted in a higher conditional R^2^, but at the cost of excluding 19 countries and 2 species from the dataset. We considered the gain in model fit did not outweigh the added spatial and taxonomic coverage. Finally, using simulated error weights and the full dataset (including censored observations), we tested how the lag period of covariates influenced fit. All lag periods offered a similar fit (marginal and conditional R^2^), and so we selected the lag most supported by the literature – 10 years, with suggestions peak population change occurs 8 years after environmental change (specifically forest loss) in mammals ^27^.

**Table S3.** Fit of nine models tested in the sensitivity analyses split across three categories: Weighting – influence of different trend observation weighting options; Censoring – impact of including different qualities of trend data; and Lag – fit under different lag periods for covariates (e.g. for a predator population monitored between 1995-2000, the Change in human density would be measured from 1995-2000, 1990-2000, and 1985-2000, respectively under the 0, 5, and 10-year lags. Fit measured as the marginal and conditional R^2^. There are varying levels of data in each model, and we summarise the frequency of countries and species this data occurs in. For the weighting models, all quantitative and qualitative-censored trends were included, with a 5-year lag on the covariates. For the censoring models, all trend observations were weighted by the simulated error, with a 5-year lag on the covariates. For the lag model, all quantitative and qualitative-censored trends were included and weighted by the simulated error.

| | Category | Marg. $R^2$ | Cond. $R^2$ | Countries (N) | Species (N) |
| --- | --- | --- | --- | --- | --- |
| Weighting | Unweighted | 0.09 | 0.25 | 75 | 50 |
|  | Weighted by trend sample size | 0.10 | 0.30 | 75 | 50 |
|  | Weighted by simulated error | 0.11 | 0.30 | 75 | 50 |
| Censoring | High quality timeseries trends | 0.11 | 0.30 | 56 | 48 |
|  | All quantitative trends | 0.10 | 0.28 | 69 | 49 |
|  | All quantitative and qualitative-censored trends | 0.11 | 0.29 | 75 | 50 |
| Lag | 0 years | 0.10 | 0.30 | 75 | 50 |
|  | 5 years | 0.11 | 0.29 | 75 | 50 |
|  | 10 years | 0.11 | 0.30 | 75 | 50 |

In our final model, we used the simulated error weights, included censored observations, and used a 10-year covariate lag. While these decisions optimized fit and data coverage, the type of weightings, the data or time lags used had little impact on inference, as model coefficients were largely similar across all parameter types (Figure S12 – S14).

**Figure S12.**
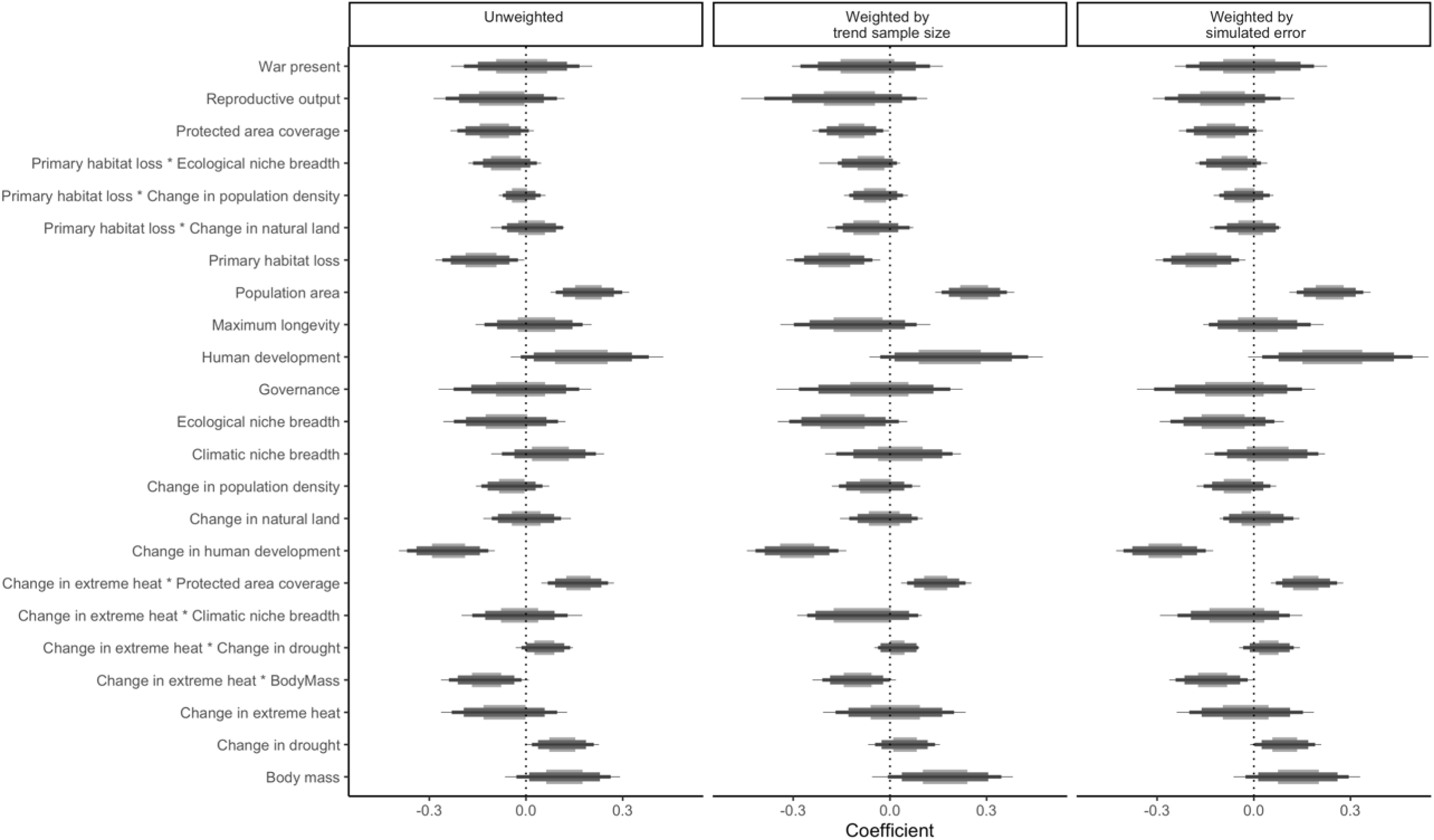
Standardized slope coefficients for the 23 main effects and interactions on the annual rate of change, comparing three models with different types of trend weighting: 1) trend values are unweighted; 2) trend values are weighted by the sample size (frequency of abundance observations used to derive trend); and 3) trend values are weighted by the simulated error. The four widths of the error bars represent different credible intervals: 50% (thickest), 80%, 95%, and 97.5% (thinnest)

**Figure S13.**
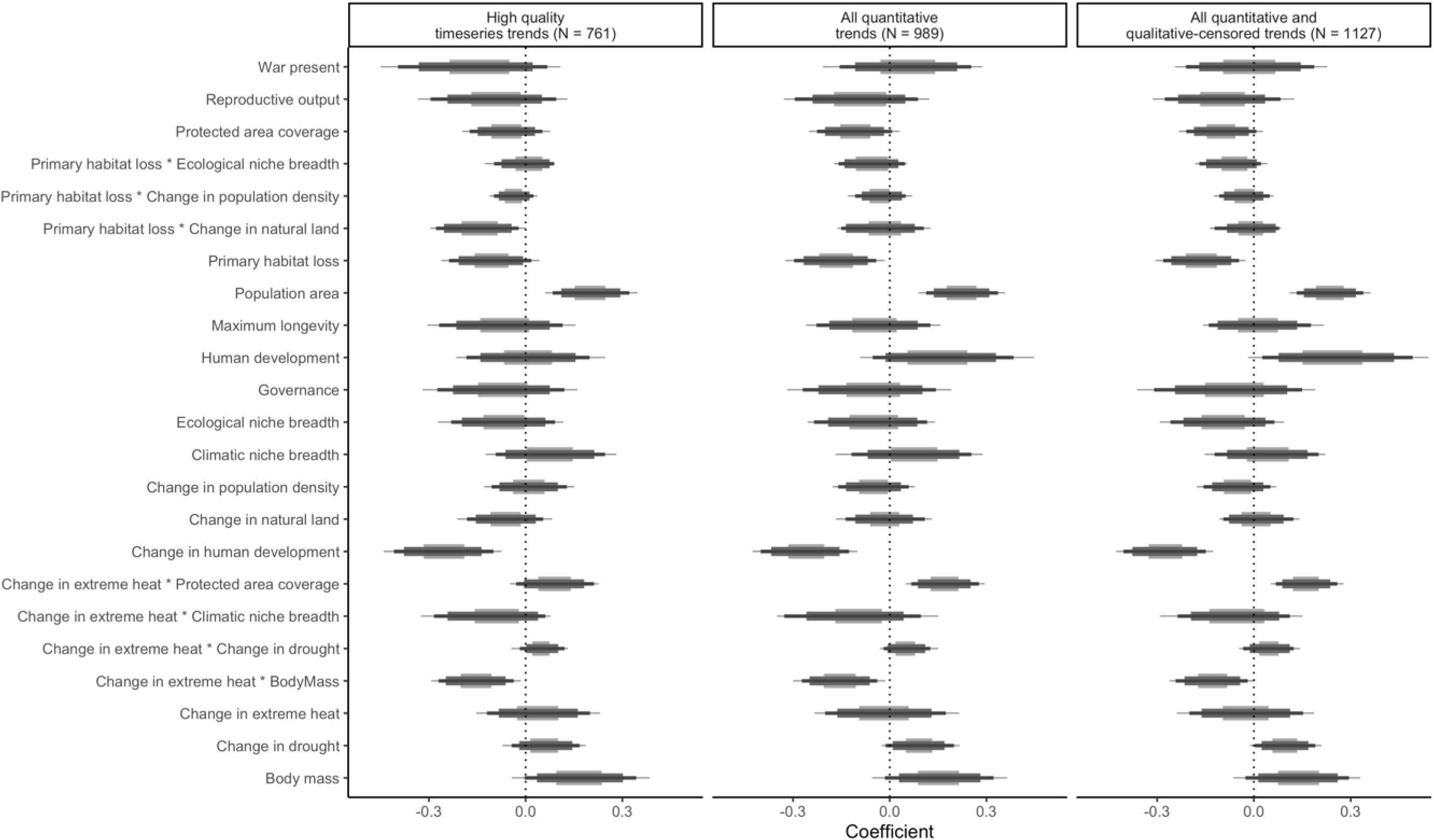
Standardized slope coefficients for the 23 main effects and interactions on the annual rate of change, comparing three models with different levels of inclusion for the trend data: 1) all timeseries trends with at least three abundance values are used; 2) all quantitative trend value are used; and 3) all trend values are used. The numbers in brackets alongside the facet titles describe the sample size of trends in the model. The four widths of the error bars represent different credible intervals: 50% (thickest), 80%, 95%, and 97.5% (thinnest)

**Figure S14.**
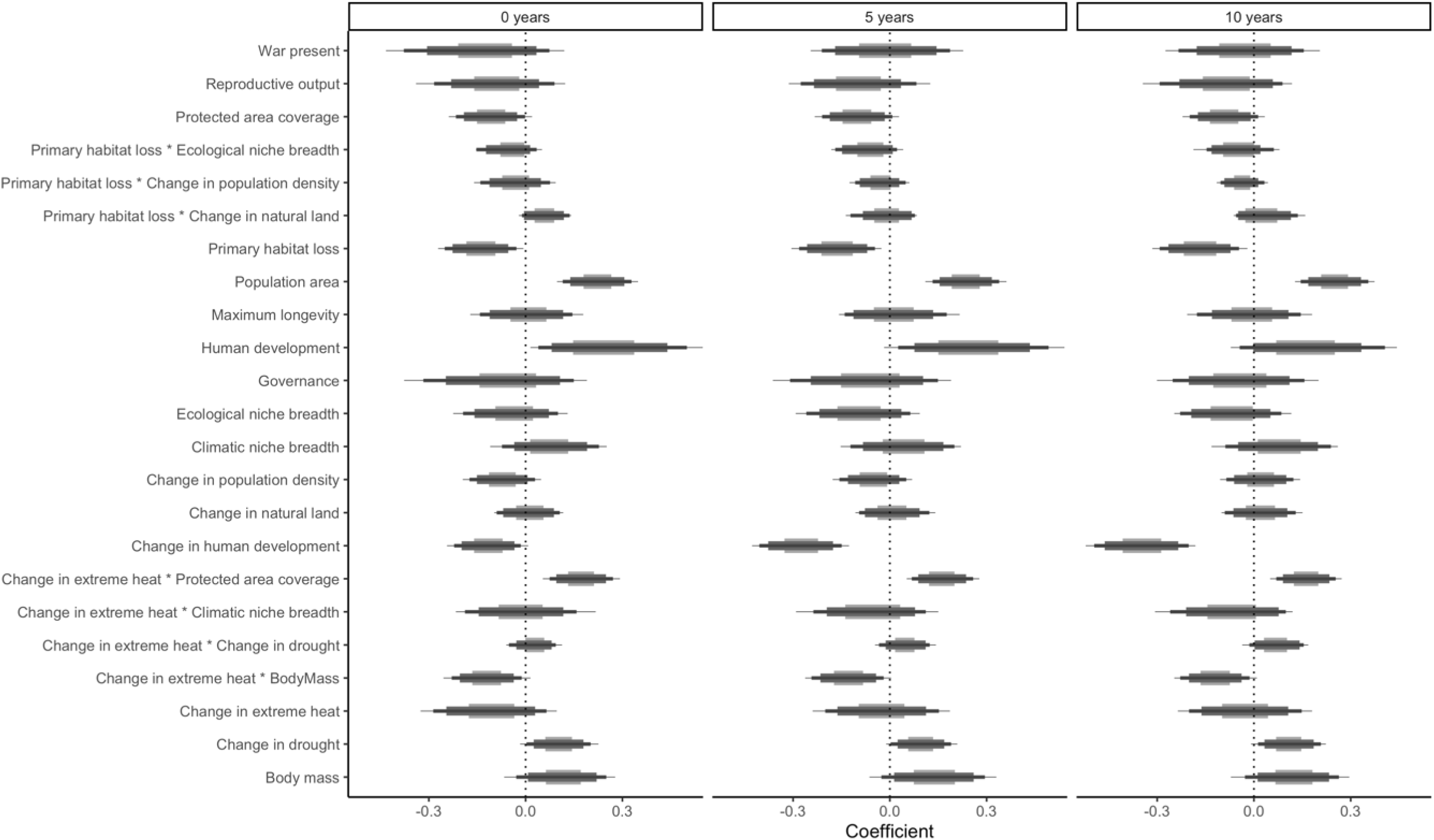
Standardized slope coefficients for the 23 main effects and interactions on the annual rate of change, comparing three models describing different lags of the model covariates: 1) 0 years; 2) 5 years; and 3) 10 years. The different lag periods only effect covariates that measure a change in the covariate over time. For example, for a predator population monitored between 1995-2000, the Change in human density would be measured from 1995-2000, 1990-2000, and 1985-2000, respectively under the 0, 5, and 10-year lags. The four widths of the error bars represent different credible intervals: 50% (thickest), 80%, 95%, and 97.5% (thinnest).

#### Parameter frequency and effects

Some parameters occurred far more frequently than others. Change in human development was the most common optional parameter, occurring in all of the selected iterations. Whilst Change in extreme heat * Protected area coverage was the most common interaction parameter (Figure S15).

**Figure S15.**
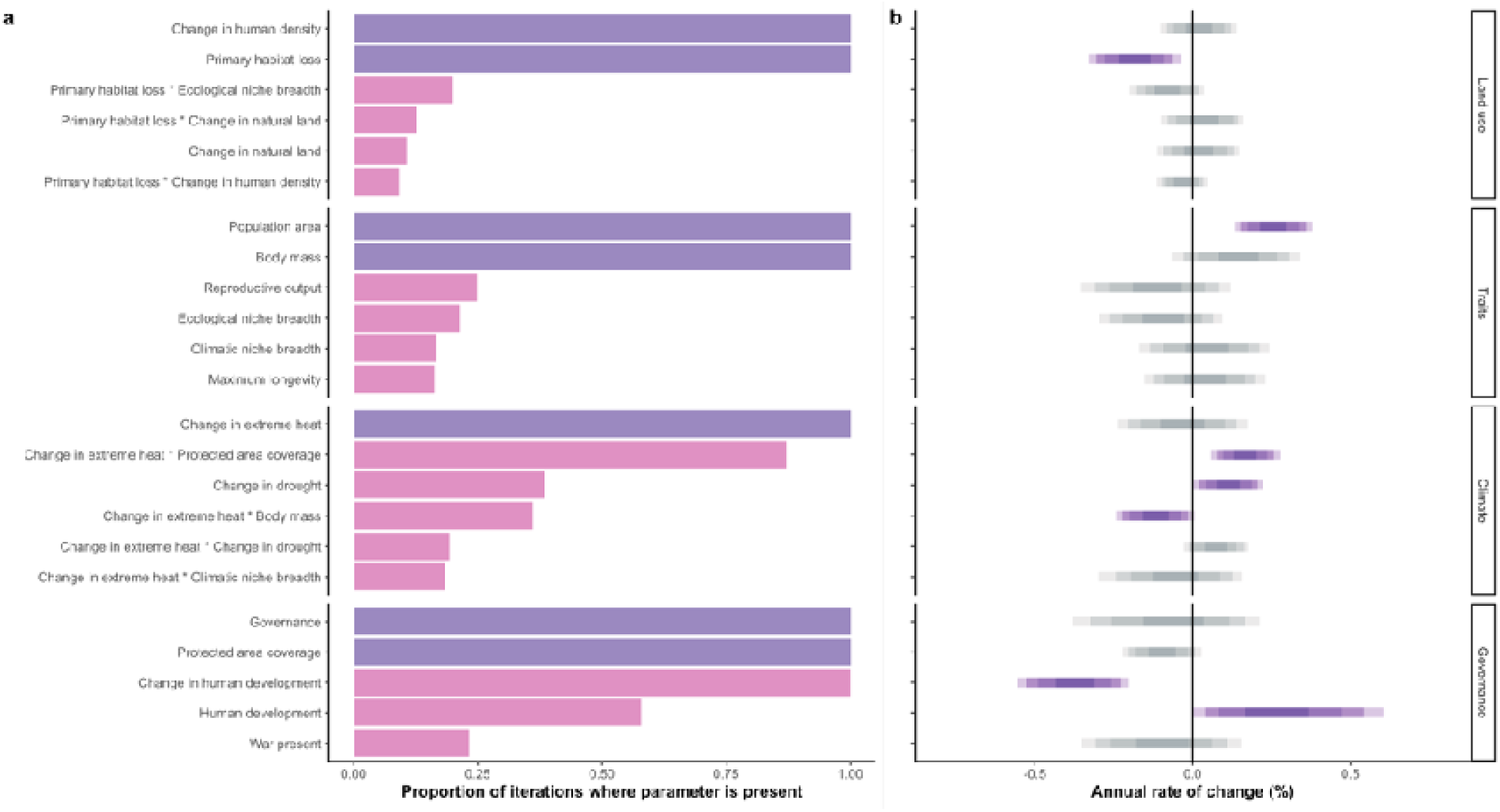
a) Proportion of iterations where parameter is present. Core parameters (purple) are present in all models, whilst optional and interaction parameters (pink) only occur in models when they are selected from Bernoulli distributions. b) Standardised coefficients of main effect and interactive model parameters. Parameters in purple have an effect at the 90% credible interval, whilst those in grey do not. The different credible interval thresholds are shown for each parameter, with darkest centre representing the 50% credible intervals, followed by the 80%, 90%, and 95% thresholds (the maximum and minimum point on each bar).

#### Model assumptions and checks

The inference model passed all standard linear mixed effect model assumptions, with residuals not showing signal of spatial or phylogenetic autocorrelation (Figure S16). However, the inference model failed to represent the more extreme observed annual rate of change (%) values, with the predicted rates of change largely ranging from -2 to 2, whilst the observed rates of change range -5 to 5; both on an inverse hyperbolic sine scale (Figure S17a). The qualitative-censored predictions largely agreed with the observed data, where censored-increasing values where primarily predicted to increase, and censored-stable values exhibited small increases and decreases (Figure S17b). The only category with reasonably poor alignment was censored-decreasing, where populations were predicted to be both increasing and decreasing. Our posterior predictive checks suggest the model produced broadly plausible values, with the independently simulated values occurring within the distribution of the observed trends, albeit failing to represent extreme values (Figure S17c). For the qualitative-censored trends, the quasi-observed values matched the simulated values almost identically (Figure S17d), which is to be expected. The inference model had a median root mean square error of 9.2%, a median marginal R^2^ of 0.15, and a median conditional R^2^ of 0.4 (Figure S18).

**Figure S16.**
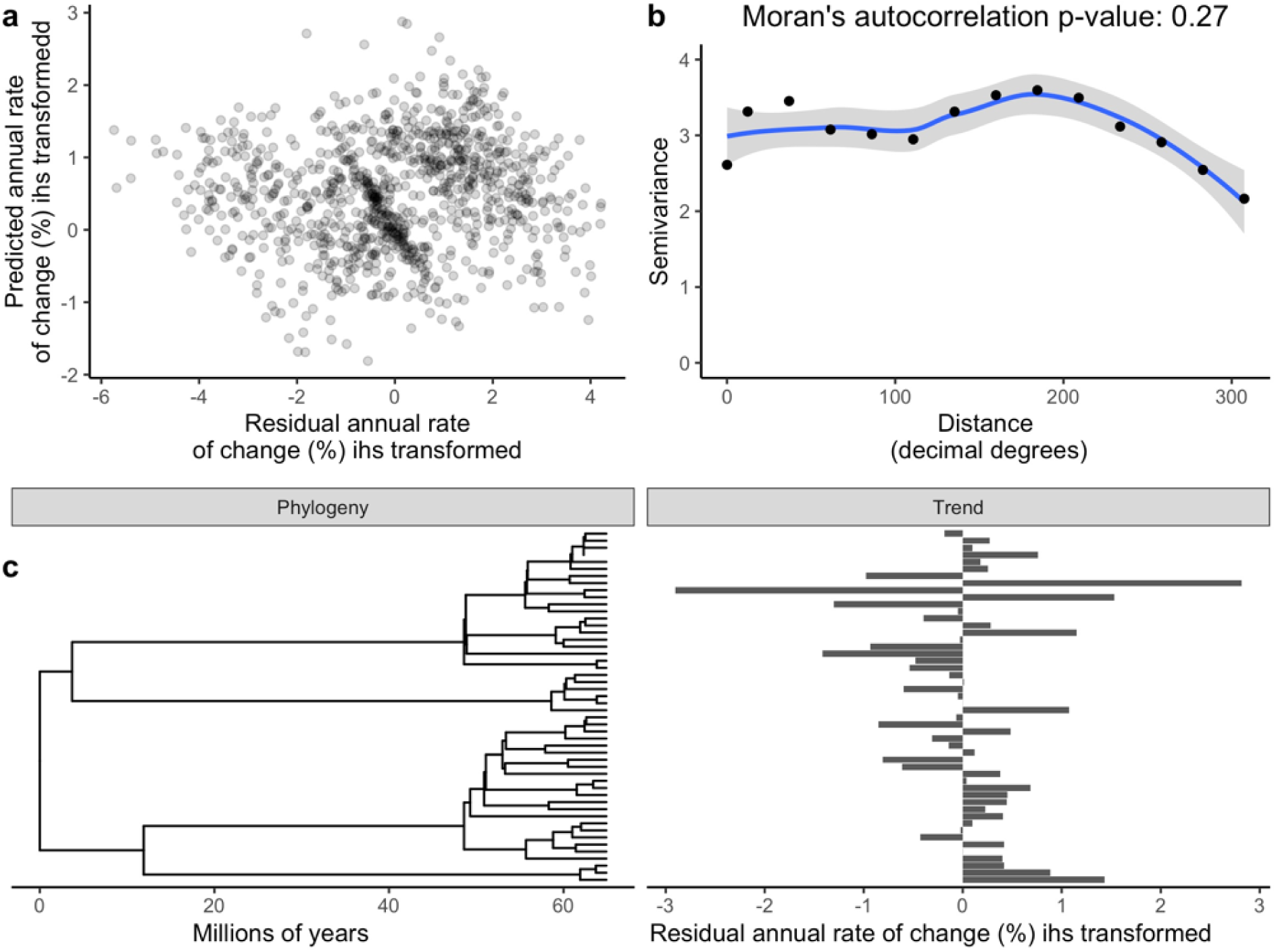
a) Median predicted annual rate of change (%) values from the inference model plotted against the median residual rates of change (%), both displayed with an inverse hyperbolic sine (ihs) transformation – the transformation used on the annual rate of change (%) within the inference model. b) Semivariance and Moran’s autocorrelation of inference model’s median residual annual rate of change (%) across distance/space (decimal degrees). c) The median residual annual rate of change (%) averaged (mean) across each species, plotted on the species’ phylogeny; with no evidence of phylogenetic autocorrelation (Pagel’s lambda = 0, p = 1). The annual rate of change (%) is ihs transformed.

**Figure S17.**
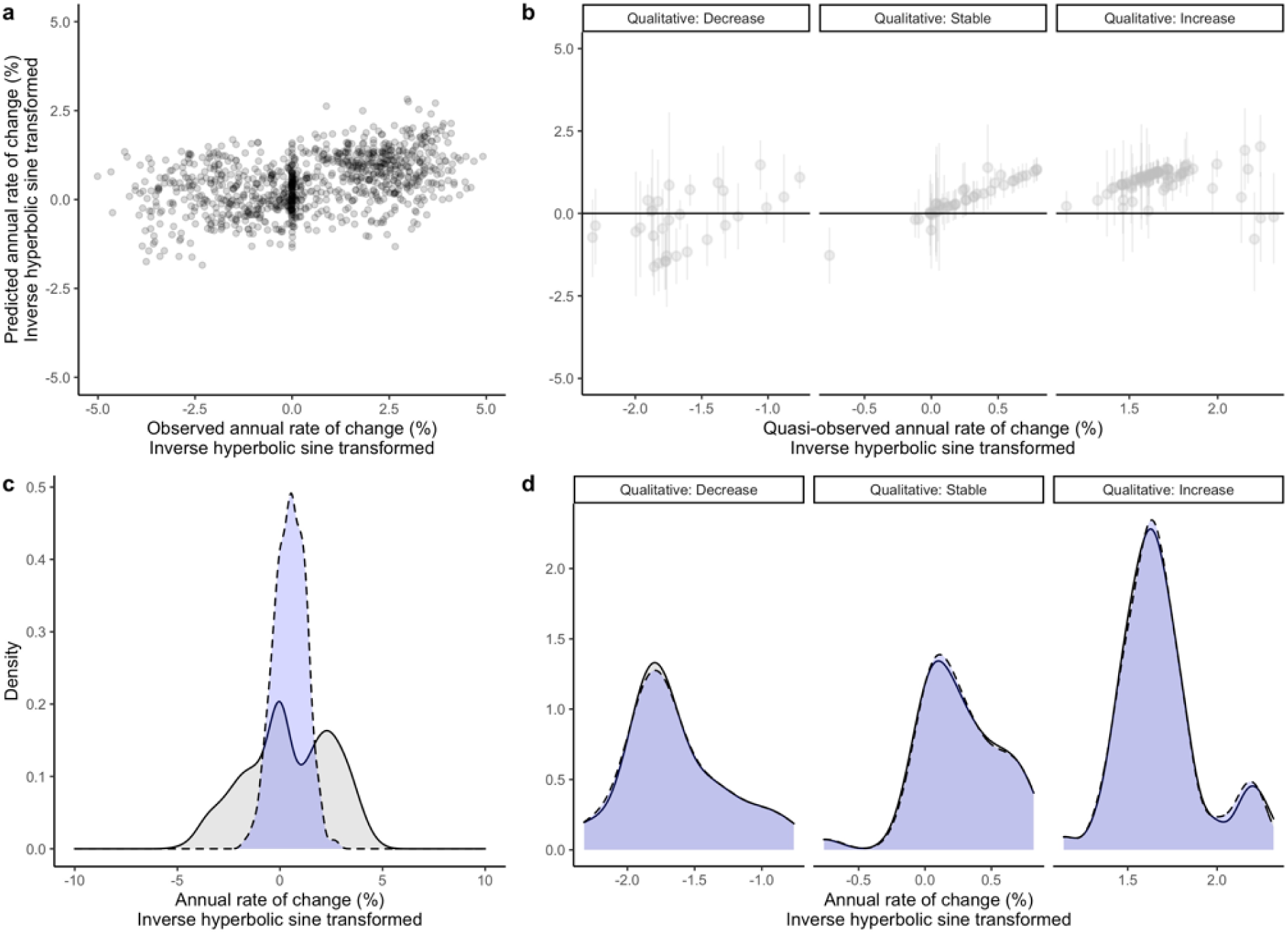
a) Median predicted annual rate of change (%) values from the inference model plotted against the observed rates of change (%), both displayed with an inverse hyperbolic sine (ihs) transformation – the transformation used on the annual rate of change (%) within the inference model. b) Median predicted annual rate of change (%) values (and 95% credible intervals) from the inference model plotted against each category of qualitative-censored values (median quasi-observed rates of change) - both displayed with an inverse hyperbolic sine (ihs) transformation. Values are quasi-observed as the true observed values are unknown. c) Distribution of observed annual rates of change (grey), compared to model simulated median annual rates of change (blue). d) Distribution of median quasi-observed annual rates of change (grey), compared to model simulated annual rates of change (blue).

**Figure S18.**
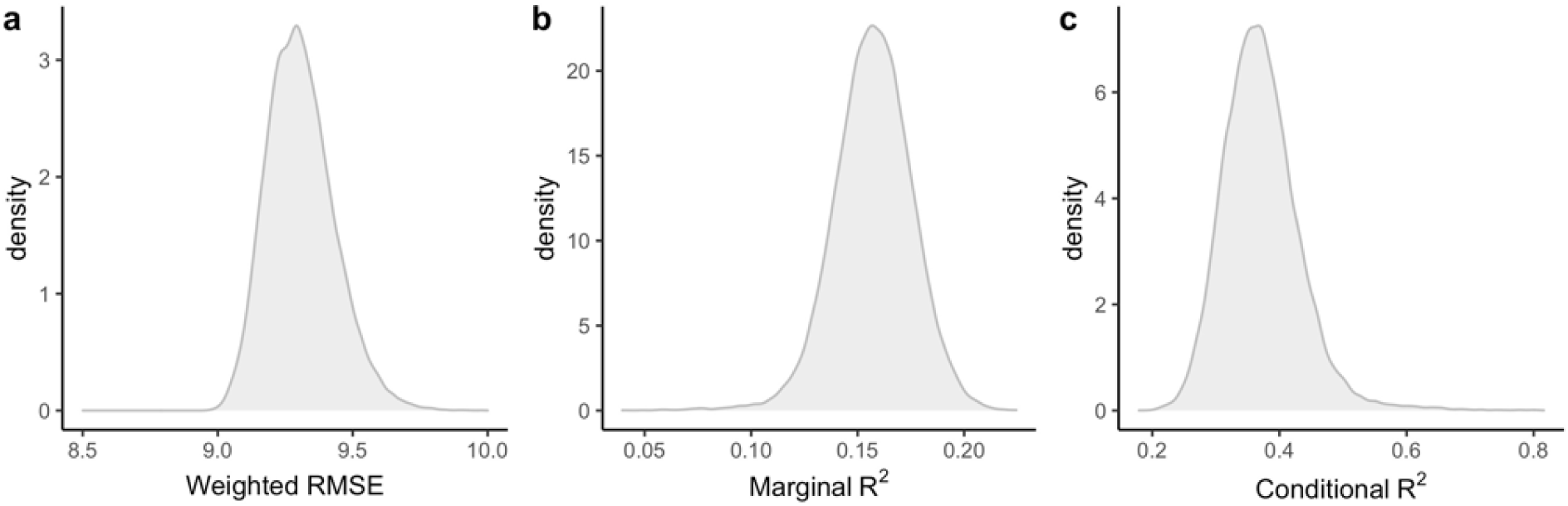
a) Distribution of the weighted root-mean-square error of the annual rate of change (%) in the inference model, comparing true to predicted values. b) Distribution of the inference model’s marginal R^2^. c) Distribution of the inference model’s conditional R^2^.

## References

1. Newbold, T. et al. Global effects of land use on local terrestrial biodiversity. Nature 520, 45–50 (2015).

2. Polaina, E., González-Suárez, M. & Revilla, E. The legacy of past human land use in current patterns of mammal distribution. Ecography (2019) doi:10.1111/ecog.04406.

3. Soroye, P., Newbold, T. & Kerr, J. Climate change contributes to widespread declines among bumble bees across continents. Science (2020) doi:10.1126/science.aax8591.

4. Spooner, F. E. B., Pearson, R. G. & Freeman, R. Rapid warming is associated with population decline among terrestrial birds and mammals globally. Glob. Change Biol. (2018) doi:10.1111/gcb.14361.

5. Trisos, C. H., Merow, C. & Pigot, A. L. The projected timing of abrupt ecological disruption from climate change. Nature (2020) doi:10.1038/s41586-020-2189-9.

6. IPBES. Summary for policymakers of the global assessment report on biodiversity and ecosystem services of the Intergovernmental Science-Policy Platform on Biodiversity and Ecosystem Services. (2019).

7. Newbold, T. et al. Has land use pushed terrestrial biodiversity beyond the planetary boundary? A global assessment. Science (2016) doi:10.1126/science.aaf2201.

8. Leung, B. et al. Clustered versus catastrophic global vertebrate declines. Nature (2020) doi:10.1038/s41586-020-2920-6.

9. van Klink, R. et al. Meta-analysis reveals declines in terrestrial but increases in freshwater insect abundances. Science (2020) doi:10.1126/science.aax9931.

10. Daskalova, G. N. et al. Landscape-scale forest loss as a catalyst of population and biodiversity change. Science (2020) doi:10.1126/science.aba1289.

11. Dornelas, M. et al. A balance of winners and losers in the Anthropocene. Ecology Letters (2019) doi:10.1111/ele.13242.

12. Hautier, Y. et al. Local loss and spatial homogenization of plant diversity reduce ecosystem multifunctionality. Nat. Ecol. Evol. (2018) doi:10.1038/s41559-017-0395-0.

13. Zavaleta, E. S., Pasari, J. R., Hulvey, K. B. & Tilman, G. D. Sustaining multiple ecosystem functions in grassland communities requires higher biodiversity. Proc. Natl. Acad. Sci. U. S. A. (2010) doi:10.1073/pnas.0906829107.

14. Fournier, A. M. V., White, E. R. & Heard, S. B. Site-selection bias and apparent population declines in long-term studies. Conserv. Biol. (2019) doi:10.1111/cobi.13371.

15. McRae, L., Deinet, S. & Freeman, R. The Diversity-Weighted Living Planet Index: Controlling for Taxonomic Bias in a Global Biodiversity Indicator. PLOS ONE 12, e0169156 (2017).

16. Humbert, J.-Y., Mills, L. S., Horne, J. S. & Dennis, B. A better way to estimate population trends. Oikos 118, 1940–1946 (2009).

17. WWF. Living Planet Report 2020 - Bending the curve of biodiversity loss. Wwf (2020).

18. Amano, T. et al. Successful conservation of global waterbird populations depends on effective governance. Nature (2018) doi:10.1038/nature25139.

19. Braga-Pereira, F., Peres, C. A., Campos-Silva, J. V., Santos, C. V. D. & Alves, R. R. N. Warfare-induced mammal population declines in Southwestern Africa are mediated by species life history, habitat type and hunter preferences. Sci. Rep. (2020) doi:10.1038/s41598-020-71501-0.

20. Clucas, B., McHugh, K. & Caro, T. Flagship species on covers of US conservation and nature magazines. Biodivers. Conserv. (2008) doi:10.1007/s10531-008-9361-0.

21. Ripple, W. J. et al. Status and ecological effects of the world’s largest carnivores. Science vol. 343 (2014).

22. Sergio, F. et al. Top predators as conservation tools: Ecological rationale, assumptions, and efficacy. Annual Review of Ecology, Evolution, and Systematics (2008) doi:10.1146/annurev.ecolsys.39.110707.173545.

23. González-Suárez, M., Lucas, P. M. & Revilla, E. Biases in comparative analyses of extinction risk: mind the gap. J. Anim. Ecol. 81, 1211–1222 (2012).

24. Johnson, T. F., Isaac, N. J. B., Paviolo, A. & González-Suárez, M. Handling missing values in trait data. Glob. Ecol. Biogeogr. (2021) doi:10.1111/geb.13185.

25. Chapron, G. et al. Recovery of large carnivores in Europe’s modern human-dominated landscapes. Science 346, 1517–1519 (2014).

26. Leclère, D. et al. Bending the curve of terrestrial biodiversity needs an integrated strategy. Nature (2020) doi:10.1038/s41586-020-2705-y.

27. Johnson, T. F., Cruz, Paula., Isaac, N. J. B., Paviolo, A. & Gonzalez-Suarez, M. CaPTrends: A global database of population trends in large terrestrial Carnivorans. Unpublished (2021).

28. WWF. Living Planet Index: Data Portal. Living Planet Index https://www.livingplanetindex.org/search (2020).

29. González-Suárez, M. & Revilla, E. Variability in life-history and ecological traits is a buffer against extinction in mammals. Ecol. Lett. 16, 242–251 (2013).

30. Pacifici, M. et al. Species’ traits influenced their response to recent climate change. Nat. Clim. Change 7, 205–208 (2017).

31. Prokosch, J., Bernitz, Z., Bernitz, H., Erni, B. & Altwegg, R. Are animals shrinking due to climate change? Temperature-mediated selection on body mass in mountain wagtails. Oecologia (2019) doi:10.1007/s00442-019-04368-2.

32. Suggitt, A. J. et al. Extinction risk from climate change is reduced by microclimatic buffering. Nature Climate Change (2018) doi:10.1038/s41558-018-0231-9.

33. Davis, K. T., Dobrowski, S. Z., Holden, Z. A., Higuera, P. E. & Abatzoglou, J. T. Microclimatic buffering in forests of the future: the role of local water balance. Ecography (2019) doi:10.1111/ecog.03836.

34. Lehikoinen, P., Santangeli, A., Jaatinen, K., Rajasärkkä, A. & Lehikoinen, A. Protected areas act as a buffer against detrimental effects of climate change—Evidence from large-scale, long-term abundance data. Glob. Change Biol. (2019) doi:10.1111/gcb.14461.

35. Fernández-Llamazares, Á., Western, D., Galvin, K. A., McElwee, P. & Cabeza, M. Historical shifts in local attitudes towards wildlife by Maasai pastoralists of the Amboseli Ecosystem (Kenya): Insights from three conservation psychology theories. Journal for Nature Conservation (2020) doi:10.1016/j.jnc.2019.125763.

36. WWF. Living Planet Report 2020 - Bending the curve of biodiversity loss. (2020).

## References

1. Johnson, T. F., Cruz, Paula., Isaac, N. J. B., Paviolo, A. & Gonzalez-Suarez, M. CaPTrends: A global database of population trends in large terrestrial Carnivorans. Unpublished (2021).

2. WWF. Living Planet Index: Data Portal. Living Planet Index https://www.livingplanetindex.org/search (2020).

3. Daskalova, G. N. et al. Landscape-scale forest loss as a catalyst of population and biodiversity change. Science (2020) doi:10.1126/science.aba1289.

4. Plummer, M. rjags: Bayesian graphical models using MCMC. R package version 3-13 (2016).

5. R Development Core Team. R Development Core Team, R: a language and environment for statistical computing. R Lang. Environ. Estat. Comput. (2020).

6. Johnson, T. F., Isaac, N. J. B., Paviolo, A. & González-Suárez, M. Handling missing values in trait data. Glob. Ecol. Biogeogr. (2021) doi:10.1111/geb.13185.

7. Goolsby, E. W., Bruggeman, J. & Ané, C. Rphylopars□: fast multivariate phylogenetic comparative methods for missing data and within-species variation. Methods Ecol. Evol. 8, 22–27 (2017).

8. Van Buuren, S. & Groothuis-Oudshoorn, K. MICE: Multivariate Imputation by Chained Equations in R. J. Stat. Softw. 10, 1–68 (2011).

9. Kuo, L. & Mallick, B. Variable selection for regression models. Sankhyā Indian J. Stat. Ser. B (1998).

## References

3. Hurtt, G. C. et al. Harmonization of global land use change and management for the period 850-2100 (LUH2) for CMIP6. Geosci. Model Dev. (2020) doi:10.5194/gmd-13-5425-2020.

4. Florczyk, A. J. et al. GHSL Data Package 2019. JRC Technical report (2019).

5. Lange, S. WFDE5 over land merged with ERA5 over the ocean (W5E5). V. 1.0. GFZ Data Services (2019).

6. Lange, S. Trend-preserving bias adjustment and statistical downscaling with ISIMIP3BASD (v1.0). Geosci. Model Dev. (2019) doi:10.5194/gmd-12-3055-2019.

7. Cucchi, M. et al. WFDE5: Bias-adjusted ERA5 reanalysis data for impact studies. Earth Syst. Sci. Data (2020) doi:10.5194/essd-12-2097-2020.

8. Thornthwaite, C. W. An Approach toward a Rational Classification of Climate. Geogr. Rev. (1948) doi:10.2307/210739.

9. Vicente Serrano, S. M., Beguiria, S. & Lopez-Moreno, J. I. A multi-scalar drought index sensitive to global warming: the standardised precipitation evapotranspiration index. J. Clim. (2010).

10. Pettersson, T. UCDP/PRIO Armed Conflict Dataset Codebook Version 19.1 . Journal of Peace Research (2019).

11. Kaufmann, D., Kraay, A. & Mastruzzi, M. The worldwide governance indicators: Methodology and analytical issues. Hague J. Rule Law (2011) doi:10.1017/S1876404511200046.

12. UNDP. Human Development Index (HDI). http://hdr.undp.org/en/content/human-development-index-hdi (2021).

13. Van Buuren, S. & Groothuis-Oudshoorn, K. MICE: Multivariate Imputation by Chained Equations in R. J. Stat. Softw. 10, 1–68 (2011).

14. UNEP-WCMC & IUCN. Protected Planet: The World Database on Protected Areas (WDPA). Available at: www.protectedplanet.net. (2021).

15. Fick, S. E. & Hijmans, R. J. WorldClim 2: new 1-km spatial resolution climate surfaces for global land areas. Int. J. Climatol. (2017) doi:10.1002/joc.5086.

16. IUCN. Habitats Classification Scheme (Version 3.1). https://www.iucnredlist.org/resources/habitat-classification-scheme (2020).

17. González-Suárez, M. Carnivora trait dataset. (2014).

18. Jones, K. E. et al. PanTHERIA: a species-level database of life history, ecology, and geography of extant and recently extinct mammals. Ecology 90, 2648–2648 (2009).

19. De MagalhÃes, J. P. & Costa, J. A database of vertebrate longevity records and their relation to other life-history traits. J. Evol. Biol. (2009) doi:10.1111/j.1420-9101.2009.01783.x.

20. Goolsby, E. W., Bruggeman, J. & Ané, C. Rphylopars□: fast multivariate phylogenetic comparative methods for missing data and within-species variation. Methods Ecol. Evol. 8, 22–27 (2017).

21. Johnson, T. F., Isaac, N. J. B., Paviolo, A. & González-Suárez, M. Handling missing values in trait data. Glob. Ecol. Biogeogr. (2021) doi:10.1111/geb.13185.

22. Nyakatura, K. & Bininda-Emonds, O. R. P. Updating the evolutionary history of Carnivora (Mammalia): A new species-level supertree complete with divergence time estimates. BMC Biol. (2012) doi:10.1186/1741-7007-10-12.

23. Daskalova, G. N. et al. Landscape-scale forest loss as a catalyst of population and biodiversity change. Science (2020) doi:10.1126/science.aba1289.

24. Plummer, M. rjags: Bayesian graphical models using MCMC. R package version 3-13 (2016).

25. R Development Core Team. R Development Core Team, R: a language and environment for statistical computing. R Lang. Environ. Estat. Comput. (2020).

26. Kuo, L. & Mallick, B. Variable selection for regression models. Sankhyā Indian J. Stat. Ser. B (1998).

27. Daskalova, G. N. et al. Landscape-scale forest loss as a catalyst of population and biodiversity change. Science (2020) doi:10.1126/science.aba1289.

